# IL-5Rα marks nasal polyp IgG4 and IgE-secreting cells in aspirin-exacerbated respiratory disease

**DOI:** 10.1101/527762

**Authors:** Kathleen M. Buchheit, Daniel F. Dwyer, Jose Ordovas-Montanes, Howard R. Katz, Erin Lewis, Juying Lai, Neil Bhattacharyya, Alex K. Shalek, Nora A. Barrett, Joshua A. Boyce, Tanya M. Laidlaw

**Affiliations:** Department of Medicine, Harvard Medical School, Boston, MA; Department of Surgery, Harvard Medical School, Boston, MA; Division of Rheumatology, Immunology, and Allergy, Brigham and Women’s Hospital, Boston, MA; Division of Otolaryngology, Brigham and Women’s Hospital, Boston, MA; Institute for Medical Engineering and Science (IMES), Department of Chemistry, and Koch Institute for Integrative Cancer Research, MIT, Cambridge, MA, USA; Broad Institute of MIT and Harvard, Cambridge, MA, USA; Ragon Institute of MGH, MIT and Harvard, Cambridge, MA, USA; Harvard-MIT Division of Health Sciences & Technology, Cambridge, MA, USA

**Author notes:** Corresponding Author: Tanya Laidlaw, MD, Address: Brigham and Women’s Hospital, 60 Fenwood Road, Building of Transformative Medicine, Rm 5002M, Boston, MA 02115.

## Abstract

**Background:** The cause of nasal polyposis in aspirin-exacerbated respiratory disease (AERD) is unknown. Elevated antibody levels have been associated with disease severity in nasal polyps, but the upstream drivers and cellular mechanisms of local antibody production in AERD remain to be investigated.

**Objective:** We sought to identify the upstream drivers and phenotypic properties of local antibody-secreting cells in nasal polyps and to understand their clinical relevance in AERD.

**Methods:** Sinus tissue was obtained from subjects with AERD, aspirin-tolerant chronic rhinosinusitis with nasal polyps (CRSwNP), aspirin-tolerant chronic rhinosinusitis without nasal polyps (CRSsNP), and healthy controls. Tissue antibody levels were quantified via ELISA and immunohistochemistry, and were correlated with clinical markers of disease severity. Tissue cytokine mRNA levels were measured with quantitative PCR (qPCR). Antibody-secreting cells were profiled with a combination of single-cell RNA-sequencing (scRNA-seq), flow cytometry and immunofluorescence.

**Results:** Tissue IgE and IgG4 were elevated in AERD compared to controls (p<0.01 for IgE and p<0.001 for IgG4, vs. CRSwNP). Total IgG and IgG4 positively correlated with the number of polyp surgeries per subject (r=0.48, p=0.011 and r=0.58, p=0.0003, respectively). Polyp IL-10 mRNA expression was higher in AERD vs. CRSwNP (p<0.05), but there were no differences in mRNA expression of type 2 cytokines. ScRNA-seq revealed increased *IL5RA*, *IGHG4*, and *IGHE* in the antibody-associated cells of subjects with AERD compared to CRSwNP. Total plasma cells and IL-5Rα^+^ plasma cell numbers in the polyp tissue from AERD exceeded those in polyps from CRSwNP (p=0.0051 and p=0.026, respectively) by flow cytometry. With immunofluorescence, we determined that IL-5Rα and IgG4 are co-expressed in antibody-secreting cells in AERD.

**Conclusions:** Our study identifies unique clusters of antibody-secreting cells in AERD defined by enrichment of transcripts encoding *IL5RA*, *IGHG4* and *IGHE*. We confirm surface expression of IL-5Rα on these cells, and identify T cells as a unique transcriptional source of IL-5. Tissue antibody levels are elevated in AERD and correlate with disease severity. Our findings suggest a role for IL-5 in facilitating local antibody production that may drive features of severe sinus disease.

**Key Messages:**

- IgG4 and IgE levels are markedly increased in nasal polyp tissue from subjects with AERD compared to aspirin-tolerant CRSwNP.
- Tissue IgG4 levels positively correlate with disease recurrence.
- IL-10 mRNA levels are significantly higher in AERD polyp tissue compared to CRSwNP tissue, but differences were not noted for type 2 cytokines or cytokines involved in class switch recombination.
- IL-5Rα transcript and protein surface expression is elevated in antibody-secreting cells from subjects with AERD and may play a role in facilitating class switching and/or survival of antibody-secreting cells.

**Capsule Summary:** Single-cell RNA-sequencing (scRNA-seq) of whole nasal polyp tissue identified increased *IL5RA*, *IGHE*, and *IGHG4* expression in the antibody-secreting cell compartment of subjects with aspirin-exacerbated respiratory disease (AERD) compared to aspirin-tolerant chronic rhinosinusitis with nasal polyps (CRSwNP). IgE and IgG4 levels are elevated in nasal polyp tissue from subjects with AERD compared to CRSwNP and correlate with disease recurrence.

## Introduction

Nasal polyps are inflammatory outgrowths of sinonasal mucosa that cause nasal obstruction and anosmia, frequently require surgical excision, and are associated with significant medical resource consumption.^1–3^ Nasal polyps are particularly severe and recurrent in aspirin-exacerbated respiratory disease (AERD), a distinct, adult-onset respiratory syndrome consisting of eosinophilic chronic rhinosinusitis with nasal polyposis (CRSwNP), asthma, and pathognomonic respiratory reactions to cyclooxygenase (COX)-1 inhibitors that involve release of multiple mast cell mediators, including tryptase, leukotriene (LT)C_4_ and prostaglandin (PG)D_2_.^4–6^ In patients with AERD, nasal polyps are frequently refractory to standard therapy and recur within two years after surgical excision in 85 percent of patients.^7^ The factors contributing to the severity and recalcitrance of the mucosal pathology in AERD remain largely unknown.

Activated B cells and antibody-secreting cells are present in nasal polyps and locally generate immunoglobulins. Subjects with recurrent nasal polyposis have elevated total nasal polyp IgA, IgG, and IgE levels.^8–11^ Potential mechanisms by which local nasal tissue immunoglobulins may contribute to nasal polyp severity include IgE- and free light chain-induced activation of polyp mast cells,^12^ IgA-enhanced eosinophil survival,^13^ and IgG-directed local complement activation.^14^ IgE antibodies to staphylococcal enterotoxins have been linked to nasal polyp pathogenesis,^15,16^ and a role for auto-antibodies in nasal polyp pathogenesis has been proposed,^8^ but no single antigen has been consistently linked to nasal polyposis in general or to AERD in particular. A previous study reported that patients with AERD have elevated serum IgG4 and slightly depressed serum IgG1 as compared to healthy controls, independent of corticosteroid exposure or IgE levels.^17^ More recently, IgG4 was identified in nasal polyp tissue from subjects with CRS and AERD and was correlated with a poor post-operative course.^18^ This suggests a possible role for IgG4 in sinus disease persistence by as yet unidentified mechanisms.

Nasal polyp tissue contains a variety of cytokines that may drive the B cell pro-inflammatory response.^19^ Type 2 cytokines, including IL-4, IL-5, IL-13, TSLP, and IL-33, as well as IL-10, are abundant in the eosinophilic nasal polyps in AERD.^6,20,21^ Some of these type 2 cytokines have been shown to influence B cell differentiation, activation and class switching, and can drive immunoglobulin production in other settings.^22,23^ In the current study, we use massively parallel single-cell RNA-sequencing (scRNA-seq) and flow cytometry to identify unique antibody-secreting cell states in nasal polyps from patients with AERD. These antibody-secreting cells express *IL5RA*, encoding for the IL-5 receptor alpha subunit (IL-5Rα), along with *IGHG4* and *IGHE*, encoding for the IgG4 and IgE heavy chains. Both IgE and IgG4 concentrations are selectively elevated in the nasal polyp tissue of subjects with AERD. Furthermore, those elevated antibody levels correlate with the severity and recurrence of the sinus disease. We suspect that the increased IgE may be pathogenic and driven in part by the effect of local T cell derived IL-5 on antibody-secreting cells in the nasal polyp tissue, and that the increased IgG4 may be a compensatory mechanism reflecting the additional influence of IL-10 from myeloid cells on local class switching. Moreover, in addition to its established role in controlling tissue eosinophilia, IL-5 may also drive pathogenic immunoglobulin production, and may be amenable to modification with IL-5-neutralizing biologic therapies.

## Methods

### Patient characterization

Subjects between the ages of 18 and 75 years were recruited from the Brigham and Women’s Hospital (Boston, MA) Allergy and Immunology clinics and Otolaryngology clinics between May 2013 and June 2018 **(Table 1)**. The local Institutional Review Board approved the study and all subjects provided written informed consent. Ethmoid sinus tissue was collected at the time of elective endoscopic sinus surgery from patients with physician-diagnosed AERD, and aspirin-tolerant CRS with and without nasal polyps with the diagnosis made based on established guidelines.^24^ Healthy control patients were undergoing sinus surgery to correct anatomic abnormalities by removal of concha bullosa. Patients were suspected of having AERD if they had asthma, nasal polyposis, and a history of respiratory reaction on ingestion of a COX-1 inhibitor, with diagnosis later confirmed in all subjects via a physician-observed graded oral challenge to aspirin. Subjects with known cystic fibrosis, allergic fungal rhinosinusitis and unilateral polyps were excluded from the study.

**Table 1.**
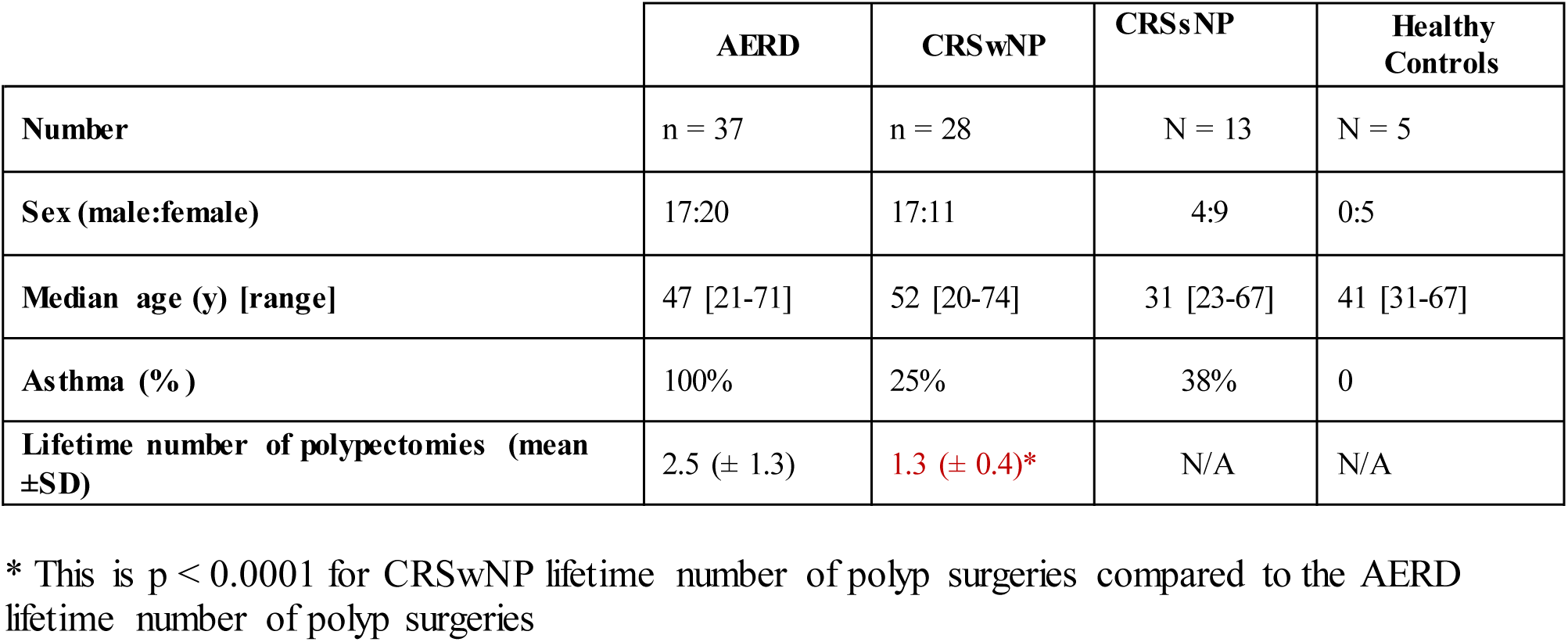
Patient characteristics.

|  | **AERD** | **CRSwNP** | **CRSsNP** | **Healthy Controls** |
| --- | --- | --- | --- | --- |
| **Number** | n = 37 | n = 28 | N = 13 | N = 5 |
| **Sex (male:female)** | 17:20 | 17:11 | 4:9 | 0:5 |
| **Median age (y) [range]** | 47 [21-71] | 52 [20-74] | 31 [23-67] | 41 [31-67] |
| **Asthma (%)** | 100% | 25% | 38% | 0 |
| **Lifetime number of polypectomies (mean ±SD)** | 2.5 (± 1.3) | 1.3 (± 0.4)* | N/A | N/A |

\* This is $p < 0.0001$ for CRSwNP lifetime number of polyp surgeries compared to the AERD lifetime number of polyp surgeries

### Polyp procurement and tissue specimen preparation

Nasal tissue was excised at the time of surgery; one tissue segment was immediately preserved in RNAlater (Qiagen, Valencia, CA) for RNA extraction, and the remaining tissue was placed in RPMI (Corning, Corning, NY) with 10% fetal bovine serum (ThermoFisher, Waltham, MA) and 1 U/mL penicillin-streptomycin for transport to the laboratory on ice. Within 2 hours of surgery, the tissue was removed from RPMI and divided into segments. One segment was transferred into Cell Lytic M Cell Lysis Reagent (Sigma-Aldrich, St Louis, MO) with 2% protease inhibitor (Roche, Indianapolis, IN) for protein extraction, and the tissue was homogenized with a gentleMACS Dissociator (Miltenyi Biotec, San Diego, CA). Supernatants were stored at −80°C. One segment was fixed in 4% paraformaldehyde, embedded in paraffin, and kept at −80°C until sectioning. For some patients, a tissue segment was also digested into a single-cell suspension for flow cytometric studies as described below.

### Tissue Digestion

Single-cell suspensions from surgical specimens were obtained using a modified version of a previously published protocol.^25^ Surgical specimens were collected into 30 mL of cold RPMI with 10% fetal bovine serum and 1 U/mL penicillin-streptomycin. Specimens were finely minced between two scalpel blades and incubated for 15 minutes at 37°C with 600 U/mL collagenase IV (Worthington, Lakewood, NJ) and 20 ug/mL DNAse 1 (Roche, Indianapolis, IN) in RPMI with 10% fetal bovine serum. After 15 minutes, samples were triturated five times using a syringe with a 16G needle and incubated for another 15 minutes. At the conclusion of the second digest period, samples were triturated an additional five times using a syringe with a 16G needle. Samples were typically fully dissociated at this step and were filtered through a 70 µm cell strainer and spun down at 500G for 10 minutes followed a rinse with ice-cold Ca/Mg free PBS (ThermoFisher, Waltham MA) to 30 mL total volume. Red blood cells were lysed using ACK buffer (ThermoFisher) for three minutes on ice to remove red blood cells, even if no red blood cell contamination was visibly seen in order to maintain consistency across patient groups. Single-cell suspensions were cryopreserved in CryoStor CS10 (Sigma) for batched flow cytometric analyses.

### Quantitative PCR

RNA was extracted from the whole nasal tissue specimens with Tri Reagent (Qiagen) and converted to cDNA by using the RT^2^ First Strand Kit (Qiagen). Expression of *IL4*, *IL5*, *IL6*, *IL7*, *IL10*, *IL13*, *IL21*, *IL23*, *TGFB1*, *IFNA1*, *CXCL12*, *CXCL13*, *PRDM1*, and *TNFSF13B* transcripts was examined using RT2 SYBR Green qPCR Master Mix (Qiagen), and normalized to glyceraldehyde-3-phosphate dehydrogenase (*GAPDH*; all primers from Qiagen).

### Immunoglobulin quantification

Protein lysate supernatants from sinonasal tissue were collected as described above. Total IgG, IgA, IgE, and IgG1, IgG2, IgG3, and IgG4 ELISAs (eBioscience, San Diego CA) were performed according to the manufacturer’s instructions. Total tissue protein levels were measured with the Pierce^®^ BCA Protein Assay kit (Thermo Scientific). Tissue immunoglobulin levels were normalized to total protein levels.

### Immunohistochemistry and Immunofluorescence

Tissue segments were fixed in 4% paraformaldehyde, embedded in paraffin, and 5 μm sections were prepared and incubated with a mouse anti-human IgG4 mAb (clone MRQ-44; Sigma) or isotype control. For immunohistochemistry, staining was developed with the EnVision System-HRP for mouse primary antibodies (Dako, Carpinteria, CA). Sections were counterstained with hematoxylin, Gill no. 2. For quantification of IgG4^+^ cells, numbers of IgG4-positive cells in photomicrographs encompassing at least 3 high power fields of subepithelial tissue were counted and expressed per high power field. For immunofluorescence, sections were blocked with 10% donkey serum, then were incubated with both mouse anti-IgG4 and a rabbit polyclonal anti-human IL-5Rα Ab (Sigma PA5-25159) or rabbit IgG in the first step, and staining was developed with AF 594 F(ab’)_2_, donkey anti-mouse IgG and AF 488 F(ab’)_2_, donkey anti-rabbit IgG.

### scRNA-Seq Analysis

Ethmoid scRNA-seq data was obtained from a previously published study,^26^ available from the dbGaP database under dbGaP accession 30434. The UMI-collapsed cells-by-genes matrix was input into Seurat,^27^ and clustering was conducted as previously described.^26^ Iterative clustering was conducted on the previously-defined Plasma cell cluster, consisting of 2,520 cells across 12 patient samples. Briefly, a list of the 1,902 most variable genes among these cells was generated by including genes with an average normalized and scaled expression value greater than 0.22 and with a dispersion (variance/mean) of between 0.22 and 7. Principal component analysis (PCA) was performed over this list of variable genes with the addition of all immunoglobulin isotype heavy chain constant regions and first 8 principal components (PCs) were selected for further analysis based on visual identification of the “elbow” in a plot of the percent variance explained per PC. Clusters were determined using FindClusters (utilizing a shared nearest neighbor (SNN) modularity optimization based clustering algorithm) on the first 8 principal components with a resolution of 0.7. Cells were then graphically displayed using Uniform Manifold Approximation and Projection (UMAP) with a minimum distance of 0.75.

To determine cytokine sources within AERD polyps, the 4,276 cells collected from AERD patient polyps were iteratively clustered in the following fashion. A list of the 1,902 most variable genes was generated using the criteria outlined above. After performing PCA, the first 15 PCs were used for clustering and UMAP display following visual inspection of the principal component elbow graph and determining the inflection point. We note that this number of PCs separated all previously-identified cell types. Cellular identities were retained from previous analysis of this dataset.^26^ Sub-analysis of the 282 myeloid cells was conducted on the 2,324 most variable genes, determined as previously mentioned. The first 5 PCs were utilized for clustering and UMAP following visual inspection of the PC elbow plot, and clustering was performed with a resolution of 0.6. Sub-analysis of the 224 T lymphocytes was conducted on the 2,587 most variable genes, determined as previously mentioned. The first 6 PCs were utilized for clustering and UMAP following visual inspection of the PC elbow plot, and clustering was performed with a resolution of 1.0.

### Flow cytometry

Cells from the digested nasal polyp single cell suspension were stained with mAbs against CD45, CD3, CD4, CD27, CD38, CD138 (eBiosciences), IL-5Rα and CD20 (BD Biosciences, Franklin Lakes NJ) to identify plasma cells/plasmablasts, B cells, and expression of the IL-5Rα. Plasma cells were defined as CD45^+^/CD3^-^/CD20^-^/CD27^+^/CD38^+^/CD138^+^ and B cells were defined as CD45^+^/CD3^-^/CD20^+^.

### Statistical analysis

Data are presented as individual points plus standard error (SEM), unless otherwise specified. For the immunoglobulin analyses, comparisons were performed with the Kruskal-Wallis one-way ANOVA due to non-Gaussian distribution of the data. Binary comparisons were carried out with the Mann-Whitney test. Significance was defined as a two-tailed *P*-value of less than 0.05. For the whole polyp mRNA cytokine analyses, comparisons were performed with an unpaired, 2-tailed *t*-test. For the IL-5Rα surface expression analysis, comparisons were carried out with the Mann-Whitney test. Linear dependence was measured with the Spearman correlation coefficient. Statistical analyses were performed using GraphPad Prism v7.0a (GraphPad Prism, La Jolla, CA).

For scRNA-seq, data was analyzed with Seurat 2.3.4^27^ implemented in RStudio. Disease-of-origin enrichment in clusters was determined in Prism using the binomial test. All violin plots, which we elected to use due to zero inflation in single-cell data, contain at minimum 292 individual data points in any one patient group. Violins were generated through default code implemented in Seurat. Statistical enrichment for genes within clusters and disease states was determined using the Tobit test for differential gene analysis.^28^ For scores in single-cell data, we report effect sizes in addition to statistical significance as an additional metric for the magnitude of the effect observed. The calculation was performed as Cohen’s d where: effect size d = (Mean1-Mean2)/(S.D. pooled).

## Results

### Study population and demographics

There were no statistically significant differences in age or sex between subjects with CRSwNP, CRSsNP, and AERD. Healthy control subjects with surgical excision of concha bullosae were all female (5 of 5 subjects). The lifetime number of polyp surgeries was significantly higher (*P* < 0.001) and the time to polyp recurrence was significantly shorter (*P* < 0.0001) in AERD subjects compared to aspirin-tolerant CRSwNP (**Table I**). All patients with AERD had physician-diagnosed asthma and their AERD diagnosis had been confirmed with an oral aspirin challenge, and 7 of 28 aspirin-tolerant CRSwNP patients had a diagnosis of asthma.

### Nasal polyp immunoglobulin levels are elevated in AERD

Polyp tissue lysates from subjects with AERD contained significantly higher concentrations of IgA, IgG, and IgE compared to sinonasal tissue from healthy controls and CRSsNP **(Figure 1 A – C)**. Polyp IgE concentrations were higher in the AERD samples than in those from CRSwNP samples (*P* <0.01), and IgA and IgG levels tended to be higher as well **(Figure 1 A – C)**. In subjects with nasal polyposis, there was a correlation between immunoglobulin levels and total lifetime number of sinus surgeries, which was used as a surrogate for disease severity **(Figure 1 D – F)**. Total immunoglobulin levels did not correlate with subject age (data not shown). There was no correlation between serum IgE and polyp IgE levels in the samples from the 22 patients for whom both serum and polyp IgE levels were available (data not shown).

**Figure 1.**
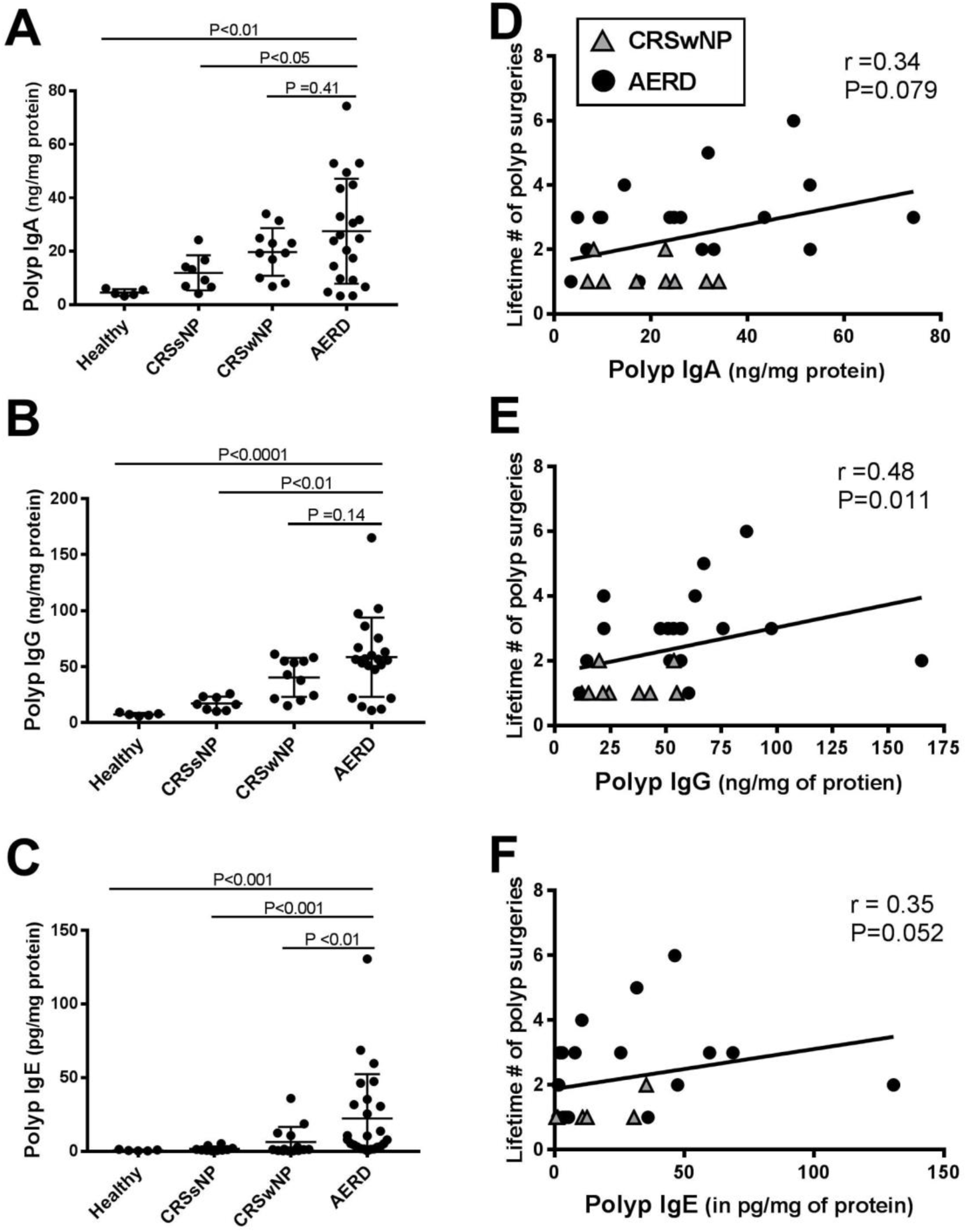
Nasal tissue antibody levels are elevated in AERD and correlate with nasal polyp recurrence. Total tissue levels of **(A)** IgA, **(B)** IgG, and **(C)** IgE were measured by ELISA from concha bullosa samples of patients without sinus inflammation (healthy controls), sinus mucosa of patients with CRSsNP, and nasal polyp tissue from patients with aspirin-tolerant CRSwNP and AERD. The nasal polyp immunoglobulin levels from patients with aspirin-tolerant CRSwNP and AERD correlate with lifetime number of polyp surgeries **(D-F)**. Data in A-C are mean ±SEM, correlation in D-F was calculated by Spearman.

To identify the isotypes responsible for the increased IgG levels in polyps from subjects with AERD, we measured specific isotypes by ELISA. Nasal polyp IgG4 protein levels were more than 6-fold higher in subjects with AERD than with CRSwNP (*P* < 0.0001), 43-fold higher in AERD compared to CRSsNP (*P* < 0.001) and close to 300-fold higher in AERD compared to healthy controls (*P* < 0.0001) **(Figure 2A)**. Furthermore, nasal polyp IgG4 levels correlated with total lifetime number of sinus surgeries (*P* < 0.01) **(Figure 2B)**. IgG4 as a percent of total IgG was significantly higher in subjects with AERD as compared to aspirin tolerant CRSwNP (*P* = 0.005) **(Figure S1)**, but there was no difference among the four phenotypic groups in their levels of IgG1, IgG2, and IgG3 as a percentage of total IgG (data not shown). Notably, nasal polyp IgG4 levels did not correlate with IgE levels in the same samples (data not shown).

**Figure 2.**
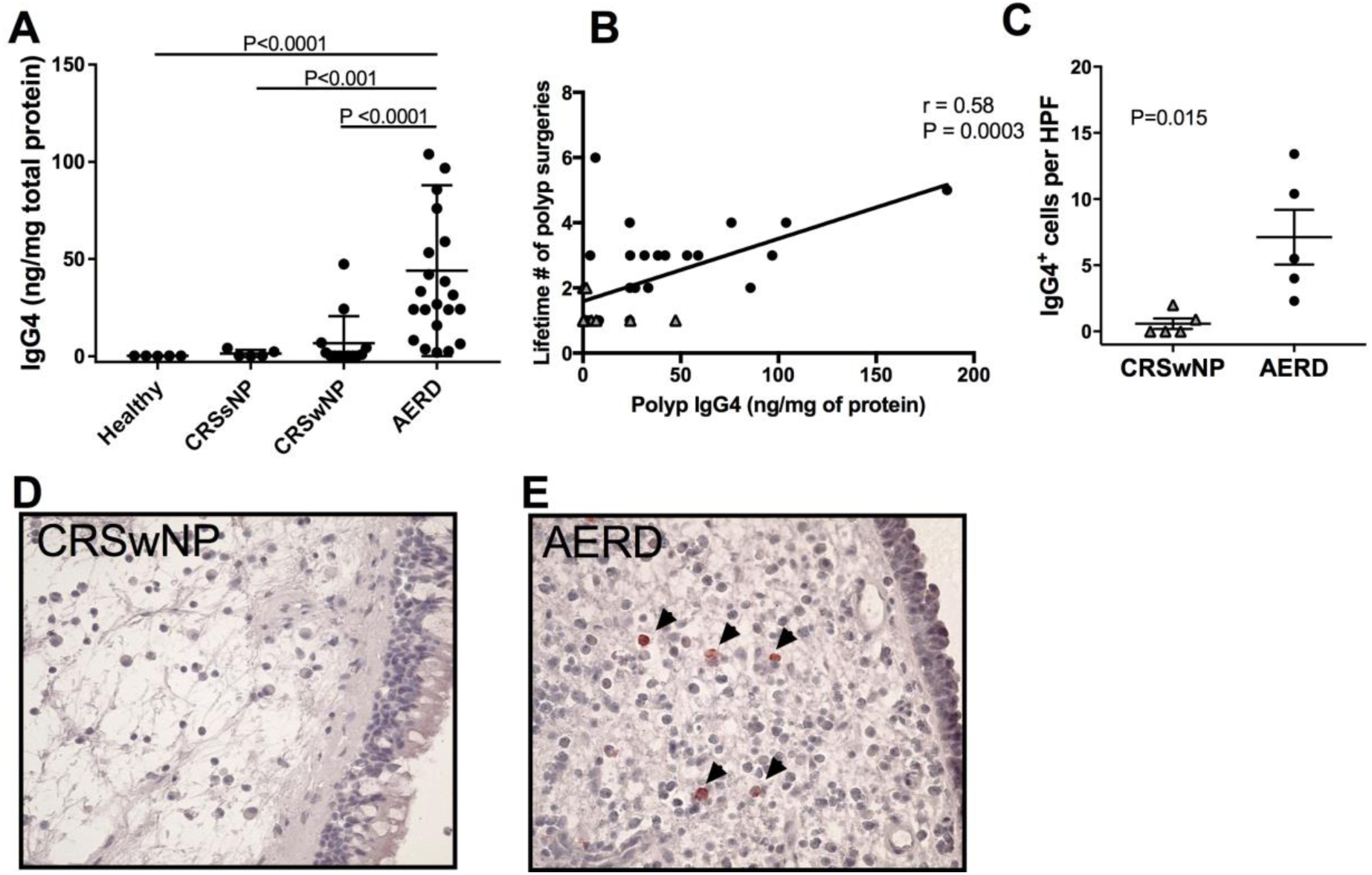
IgG4 isotype and IgG4^+^ antibody-secreting cells are specifically elevated in nasal polyps from patients with AERD. **(A)** IgG4 levels measured in tissue lysates from healthy patients without sinus inflammation, sinus mucosa of patients with CRSsNP, and nasal polyp tissue from patients with aspirin-tolerant CRSwNP and AERD. **(B)** Nasal polyp IgG4 levels from patients with aspirin-tolerant CRSwNP and AERD correlate with lifetime number of polyp surgeries. **(C**) Number of IgG4^+^ lymphocytes per HPF from nasal polyp tissue of patients with aspirin-tolerant CRSwNP and AERD, n=5 for each group. Data in A and C are mean ±SEM. **(D,E)** Representative samples (CRSwNP, **D** and AERD, **E**) of nasal polyp tissue stained with anti-IgG4. Black arrows identify IgG4^+^ cells.

To further confirm our findings, we immunohistochemically evaluated nasal polyp tissue for IgG4^+^ antibody-associated cells. We found that subjects with AERD had over 5-fold more IgG4^+^ cells compared to subjects with CRSwNP **(Figure 2C-E)**. The IgG4^+^ cells did not appear to organize into lymphoid aggregates.

### Type 2 cytokine and B cell function-related mRNA expression in nasal polyp subsets

To determine the factors driving local IgG4 production in the nasal polyp tissue of subjects with AERD, we used qPCR to measure mRNA for a number of cytokines potentially involved in immunoglobulin production and class switch recombination in the nasal polyp tissue of subjects with AERD and CRSwNP. There was significantly more *IL10* mRNA present in the whole nasal polyp tissue of subjects with AERD compared to CRSwNP (*P* = 0.034) **(Table 2)**, but no differences in type 2 cytokine mRNA levels measured, including *IL4* and *IL13*, or in other cytokines or growth factors relevant to B cell function, including *IL6* and *TGFB1* **(Table 2)**. We could not detect *IL21* transcript in a sufficient number of samples to make a comparison between groups (data not shown). IL-5 protein was below the limit of ELISA detection in most of our samples (data not shown).

**Table 2:** Whole polyp qPCR for Type 2 cytokine and B cell-related genes. Expression of cytokine transcripts in whole nasal polyp tissue specimens is shown for a panel of B cell active cytokines, normalized to glyceraldehyde-3-phosphate dehydrogenase; unpaired, 2-tailed *t*-test.

|  | AERD | Number | CRSwNP | Number | P-value |
| --- | --- | --- | --- | --- | --- |
| *IL4* | 0.0016 ± 0.00075 | 13 | 0.00037± 8.526e-005 | 9 | 0.20 |
| *IL5* | 0.088 ± 0.047 | 8 | 0.0065 ± 0.0026 | 8 | 0.10 |
| *IL6* | 0.079 ± 0.032 | 8 | 0.0069 ± 0.0028 | 8 | 0.07 |
| *IL7* | 0.0090 ± 0.015 | 8 | 0.0060 ± 0.0064 | 8 | 0.65 |
| *IL10* | **0.0017 ± 0.00045** | 13 | **0.00047 ± 0.00012** | 9 | **0.037** |
| *IL13* | 0.0010 ± 0.00052 | 13 | 0.00094 ± 0.00060 | 9 | 0.91 |
| *IL23* | 0.065 ± 0.087 | 8 | 0.018 ± 0.025 | 8 | 0.23 |
| *TGFB1* | 0.025 ± 0.017 | 8 | 0.025 ± 0.012 | 8 | 0.87 |
| *IFNA1* | 0.0013 ± 0.00078 | 8 | 0.0015 ± 0.0024 | 8 | 0.84 |
| *CXCL12* | 0.36 ± 0.0070 | 9 | 0.031 ± 0.0083 | 7 | 0.65 |
| *CXCL13* | 0.0015 ± 0.00046 | 9 | 0.0046 | 7 | 0.27 |
| *PRDM1* | 0.012 ± 0.0028 | 9 | 0.020 ± 0.0084 | 7 | 0.31 |
| *TNFSF13B* | 0.0017 ± 0.0017 | 9 | 0.0028 ± 0.00082 | 7 | 0.17 |

### ScRNA-seq identifies a transcriptionally distinct antibody-producing cell cluster unique to AERD

To extend our primary observations and identify the cellular sources of class switch-associated cytokines in an unbiased fashion, we utilized a previously-generated scRNA-Seq dataset of surgically-resected and dissociated nasal polyp tissue from a cohort of three subjects with AERD, three subjects with aspirin-tolerant CRSwNP, and five subjects with CRSsNP, specifically focusing on the previously-identified antibody-secreting cell clusters.^26^ Iterative clustering of these populations yielded 9 clusters (**Figure 3A**), all of which contained cells derived from at least eight donors and all three disease states (**Figure S2**). The majority of cluster-defining genes encoded immunoglobulin components (**Supplementary Table E1**). As previously observed,^26^ kappa and lambda light chain usage underlies a major division between clusters (**Figure S3A**). Little *IGHM* or *IGHD* expression was observed (**Figure S3B),** while robust expression of IgA and IgG isotype regions informs the remaining clusters, indicating that the majority of antibody-secreting cells detected were class-switched (**Figure S2C**).

**Figure 3.**
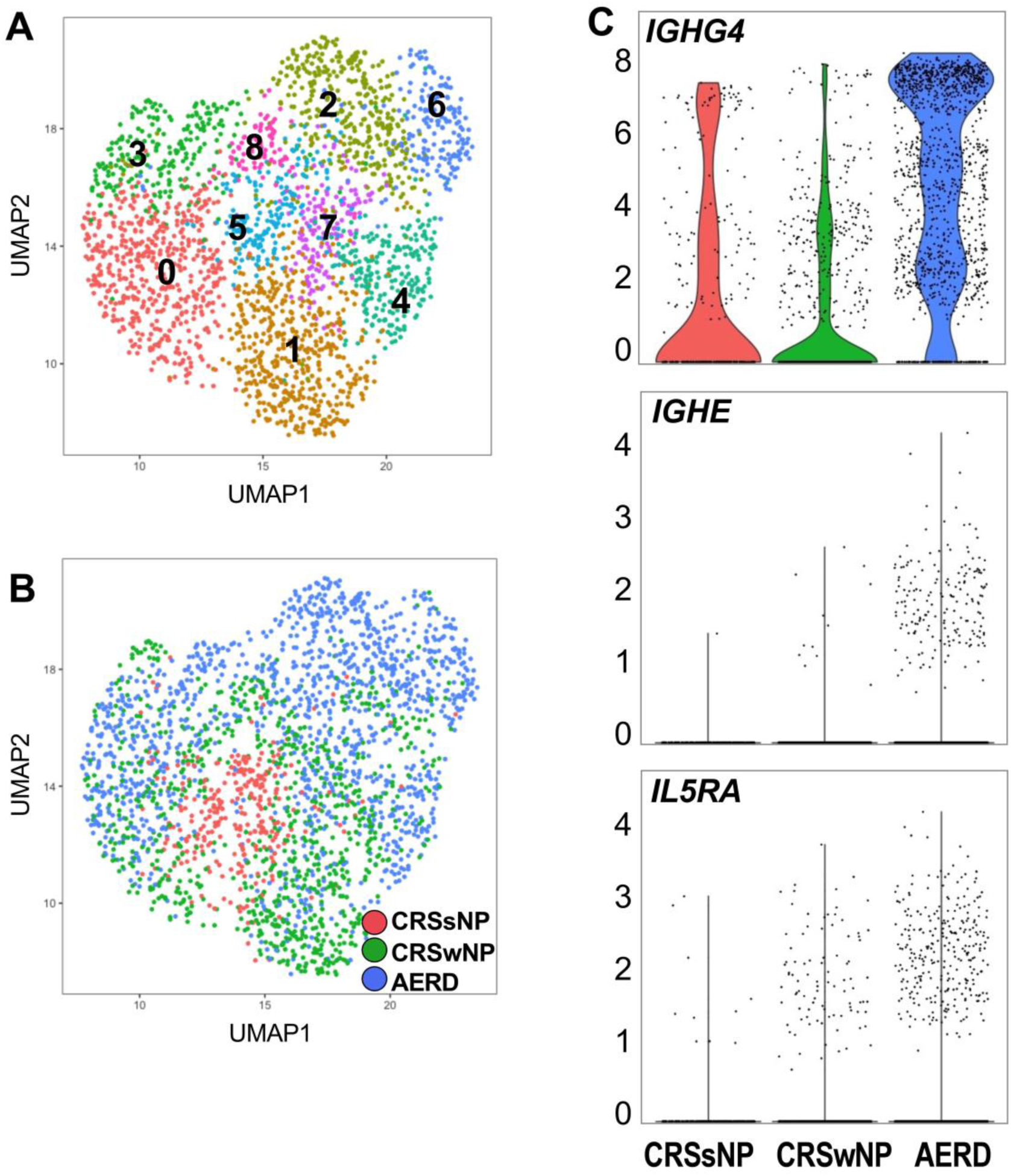
ScRNA-Seq of antibody secreting-cell populations from sinus tissue of subjects with CRSsNP (n=5), CRSwNP (n=3) and AERD (n=3). **A.** UMAP plot of 2,520 antibody-secreting cells from sinonasal tissue of CRSsNP, CRSwNP and AERD patients, indicating 9 clusters identified through a shared nearest neighbor analysis. **B.** UMAP plot of sinonasal antibody-secreting cells, colored by disease of origin. Statistical enrichment for AERD disease-of-origin was observed for cluster 2 (p<1×10^−15^), cluster 3 (p<1×10^−15^), cluster 4 (p<2×10^−12^), cluster 6 (p<1×10^−15^), and cluster 7 (p<1×10^−15^) **C.** Violin plots of select genes significantly enriched in AERD relative to CRSsNP and CRSwNP within sinonasal antibody-secreting cell populations, including *IGHG4* (p<1×10^−201^), *IGHE* (p<2×10^−34^), and *IL5RA* (p<2×10^−21^). Cohen’s d effect size for AERD relative to CRSwNP is 1.57, 0.50, and 0.37, respectively for the 3 transcripts.

Interestingly, clusters 2, 3, 4, 6, and 7 showed significant enrichment for cells derived from AERD patients (**Figure 3B**).

To understand the disease-specific differences underlying our clustering, we specifically compared transcript expression between AERD, CRSwNP and CRSsNP-derived antibody-secreting cells (**Supplementary Table E2**). *IGHG4*, encoding the IgG4 constant region, was significantly increased in AERD relative to CRSwNP and CRSsNP (**Figure 3C**, **Figure S3C**), confirming a local source for the increased protein levels **(Fig. 2A)**. We similarly saw enriched expression for *IGHE*, encoding the IgE constant region (**Figure 3C, Figure S3D**).

To gain additional insights into potential mechanisms regulating these AERD-enriched antibody-secreting cell clusters, we further analyzed the underlying gene lists to look for unique cell-surface receptor expression. Despite not identifying significant differences in *IL5* mRNA levels in bulk tissue, we found that all AERD-derived antibody-secreting cells were significantly enriched for *IL5RA* (**Figure 3C, Figure S3E**), encoding IL-5Rα, and further observed that this was the sole enriched cytokine receptor **(Supplemental Table E2).**

To understand the contribution of different cell types to cytokine production in AERD, we utilized our previously-generated scRNA-seq dataset of dissociated nasal polyp cells from subjects with AERD. ScRNA-seq of all polyp cells revealed the cellular identity of respiratory epithelial, stromal and immune cell types in the nasal polyp tissue (**Figure S4**). We examined the transcripts of each cell type to identify the potential cell-of-origin for type 2 cytokines possibly involved in class switching to IgE and IgG4 in AERD. Myeloid cells were the dominant source of *IL10*, with IL-10 expression specifically mapping to the previously identified *S100A8*-expressing inflammatory DC-3 and *C1Q*-expressing macrophages^26^ within the myeloid cluster (**Figure S4 B**). IL-5 expression was restricted to the T cell cluster, and sub-analysis indicated that these T cells co-expressed *IL13* and *HPGDS*, suggestive of the recently identified Th2A cell^29^ (**Figure S4 C**).

### Surface expression of IL-5Rα antibody secreting cells from nasal polyps

To further evaluate differences in antibody-secreting cells between subjects with AERD and CRSwNP, we examined plasma cells in the nasal polyp single cell suspensions from subjects with AERD and CRSwNP. We flow cytometrically quantified plasma cells as CD45^+^/CD3^-^/CD20^-^/CD27^+^/CD38^+^/CD138^+^ and found that subjects with AERD have significantly higher numbers of plasma cells within their nasal polyps compared to tissue from subjects with aspirin-tolerant CRSwNP (P = 0.0051) **(Figure 4A)**. There was no significant difference in the percentage of CD45^+^ cells that were B cells in subjects with AERD (5.5 ± 1.8%) vs. CRSwNP (3.5 ± 0.7%, P = 0.35). The plasma cells in nasal polyps from subjects with AERD also had greater surface expression of IL-5Rα compared to tissue from subjects with aspirin-tolerant CRSwNP (P = 0.026) **(Figure 4B, C)**. Using immunofluorescence, we examined nasal polyp tissue from 3 patients with AERD and identified co-localization of IL-5Rα and IgG4 in antibody-secreting cells (**Figure 4D,** representative sample).

**Figure 4.**
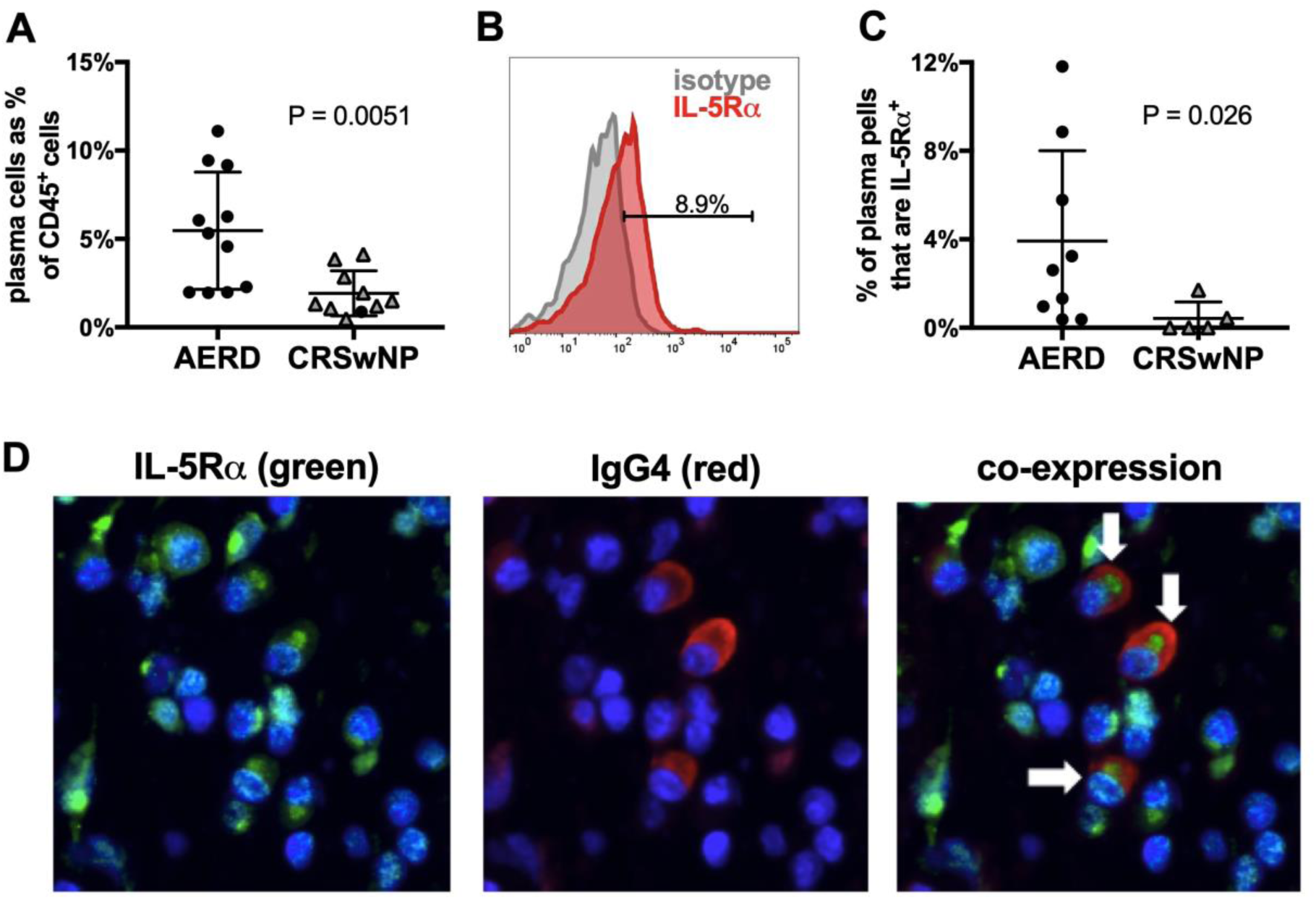
Flow cytometric characterization of plasma cells and immunofluorescence of polyp tissue. **(A)** Nasal polyp plasma cell frequency as a percentage of CD45^+^ cells, **(B)** representative histogram of IL-5Rα surface expression (with isotype positivity subtracted) on plasma cells from an AERD patient, **(C)** plasma cell surface expression of IL-5Rα, and **(D)** immunofluorescence staining of nasal polyp tissue for plasma cells (white arrows) co-expressing IL-5Rα (green) and IgG4 (red).

## Discussion

Neither the regulatory factors nor the direct consequences of local immunoglobulin production in nasal polyp tissue are known. Furthermore, differences in immunoglobulin production levels between subjects with aspirin-tolerant CRSwNP and AERD had not previously been recognized. Due to the potential importance of IgE and IgG4 in AERD pathogenesis and the potential for additional antibody-driven effector mechanisms, we sought to characterize local immunoglobulin production in nasal polyp tissue in subjects with AERD and identify factors that influence the relevant antibody-secreting cells.

We tested whole nasal polyp extracts from patients with AERD, aspirin-tolerant CRSwNP, and controls with CRSsNP and concha bullosa tissue (as a surrogate healthy control tissue) for concentrations of discrete antibody isotypes. As anticipated, polyps contained all antibody isotypes at higher concentrations than in controls, and total polyp IgG (r=0.48, P=0.011) and IgG4 (r=0.58, P=0.0003) correlated with lifetime numbers of polyp surgeries, with a weak IgE correlation as well (r=0.35, P=0.052) **(Figure 1E-F and Figure 2 B)**. While all antibody levels tended to be higher in the polyps from AERD subjects than those from CRSwNP, the differences between these two groups in total IgE **(Figure 1C)** and IgG4 **(Figure 2)** were remarkable. Moreover, polyp IgE levels did not correlate with serum IgE from the same subjects (data not shown), suggesting that IgE was synthesized locally. IgE-producing cells are notoriously difficult to detect due to very low receptor density compared with other isotypes, and their ephemeral nature in memory B cell pool of blood and secondary lymphoid organs.^30^ However, IgG4^+^ cells were readily detectable in the AERD polyps, and far more numerous than in the aspirin-tolerant control polyps, and rare IgE-expressing cells could be observed through scRNA-seq analysis (**Figures 2C-E, 3, S3**). These observations support mechanisms that specifically regulate the local productions of IgE and IgG4 in nasal polyps, and that strongly differentiate AERD from CRSwNP. It is suspected that local tissue mast cell activation contributes to nasal tissue inflammation in AERD, though the underlying mechanisms that lead to chronic mast cell activation in the tissue have not been elucidated. Although many subjects with AERD lack classic atopy,^31^ they do tend to have elevated systemic IgE levels.^31^ A recent study reported that treatment with omalizumab, a monoclonal antibody against IgE, improved sinonasal symptoms in patients with AERD and also decreased urinary PGD_2_ metabolite and leukotriene E_4_ levels, both of which are likely derived from mast cells, by ~90%.^32^ Therefore, the elevated levels of IgE **(Figure 1C)** we have found in the local nasal tissue could be instrumental to the mast cell activation within the nasal polyp tissue in AERD. However, our data suggests that the IgG4 production more strongly contributes to the severity and aggressive regrowth of nasal polyps observed in these patients.

Whereas locally-generated IgE may permit mast cells, basophils, and other FcεRI-bearing effector cells to respond to cryptic or microbial antigens, the pathophysiologic significance of IgG4 is not clear. Like IgE production, IgG4 production by B cells is regulated by IL-4/IL-13 signaling, but the balance toward IgG4 is controlled by the regulatory cytokine IL-10.^33^ ScRNA-seq analysis of nasal polyp cells from subjects with AERD revealed expression of *IL10* by macrophages and inflammatory DC3, with a minor contribution from the T cell compartment **(Figure S4**). Our finding that AERD polyps express more than three-fold higher levels of *IL10* mRNA (but not other B cell active cytokines) than CRSwNP tissue **(Table 2)** is consistent with regulatory T cells or myeloid cells driving IgG4 production in response to chronic antigen exposure. IgG4 may have an immunoregulatory role in patients with allergic sensitization^34^ and is involved in the immune response to invasive parasites.^35^ However, it is also elevated in pathologic conditions including eosinophilic esophagitis^36^ and IgG4-related diseases, a group of fibro-inflammatory disorders involving multiple organ systems.^37^ Given that IgG4 can potentially block antigen binding to IgE, it is also possible that it could modify skin test reactivity in patients with AERD, who are frequently non-atopic, as it may in subjects with eosinophilic esophagitis who respond clinically to food protein withdrawal even without evidence for IgE sensitization.

We then sought to identify cell type-intrinsic factors that might favor the production of IgE and IgG4 over other isotypes in AERD polyps. Massively parallel scRNA-Seq can reveal cell-type and disease-specific differences in mRNA expression profiles by revealing the most strongly differentially expressed transcripts. Accordingly, we identified distinct clusters of antibody-secreting cells in AERD notable for their strong expressions of *IGHG4* and *IGHE* and also distinguished by *IL5RA* expression **(Figure 3B and 3C)**. We verified that AERD polyps contained substantially greater numbers and percentages of plasma cells bearing surface IL-5Rα than did CRSwNP control polyps (**Figure 4C**), and we found that nasal polyp antibody-secreting cells in subjects with AERD express both IL-5Rα and IgG4 (**Figure 4D**). Though best known for its survival-sustaining effects on eosinophils, IL-5 was originally described as a factor required for the activation, proliferation, and differentiation of mouse B cells into antibody-secreting plasma cells,^38–40^ and acts as a strong survival factor for mouse plasma cells.^41^ Further, IL-5 has been shown to act synergistically with IL-4 to increase lymphocyte production of IgE from human lymphocytes *in vitro*^42^ and IL-5 production is known to be associated with IgE levels in humans *in vivo*.^43^ We suspect that IL-5 signals selectively to plasma cells that generate both IgE and IgG4, potentially sustaining their survival and promoting antibody production. What selectively regulates the expression of IL-5Rα on IgG4 and IgE-secreting cells will be of great interest to determine.

Humanized monoclonal antibodies against IL-5 and IL-5Rα show efficacy in the treatment of eosinophilic asthma and nasal polyposis.^44,45^ A Phase 2 trial of IL-5 inhibition with mepolizumab in patients with nasal polyposis showed a therapeutic effect with a reduction in both polyp size and patient symptoms,^46^ and we recently demonstrated that mepolizumab improved upper respiratory symptoms and asthma control in subjects with AERD.^47^ However, another recent study of dexpramipexole, an experimental drug that depletes nearly all eosinophils from within the nasal polyp tissue, failed to show any symptomatic improvement or any reduction in nasal polyp size.^48^ Taken together with our current findings, we suspect that IL-5-targeting monoclonal antibodies may alter the survival and function of IL-5Rα^+^ antibody-secreting cells, which may contribute to the mechanism of their therapeutic benefit.

---

**Supplementary Figure 1.**
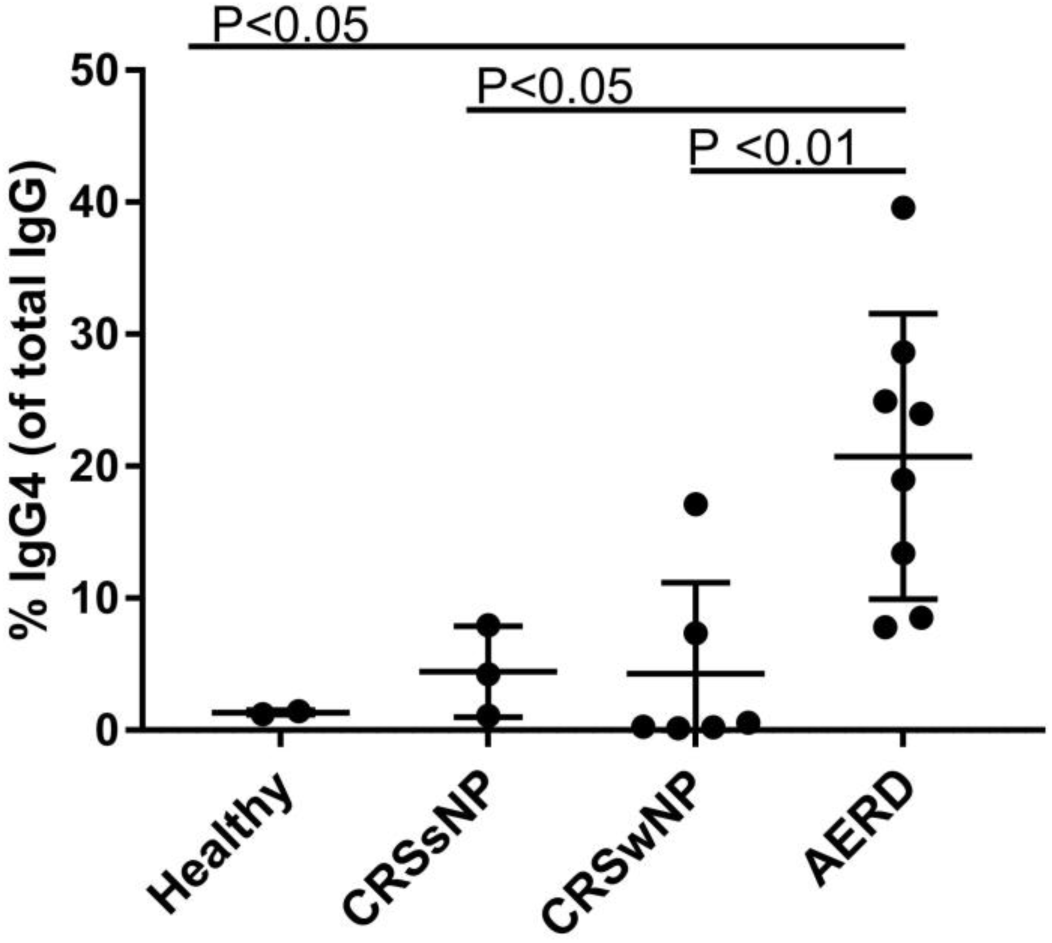
IgG4 levels measured in tissue lysates as a percent of total IgG tissue lysate levels from healthy patients without sinus inflammation, sinus mucosa of patients with CRSsNP, and nasal polyp tissue from patients with aspirin-tolerant CRSwNP and AERD. Data are mean ±SEM.

**Supplementary Figure 2.**
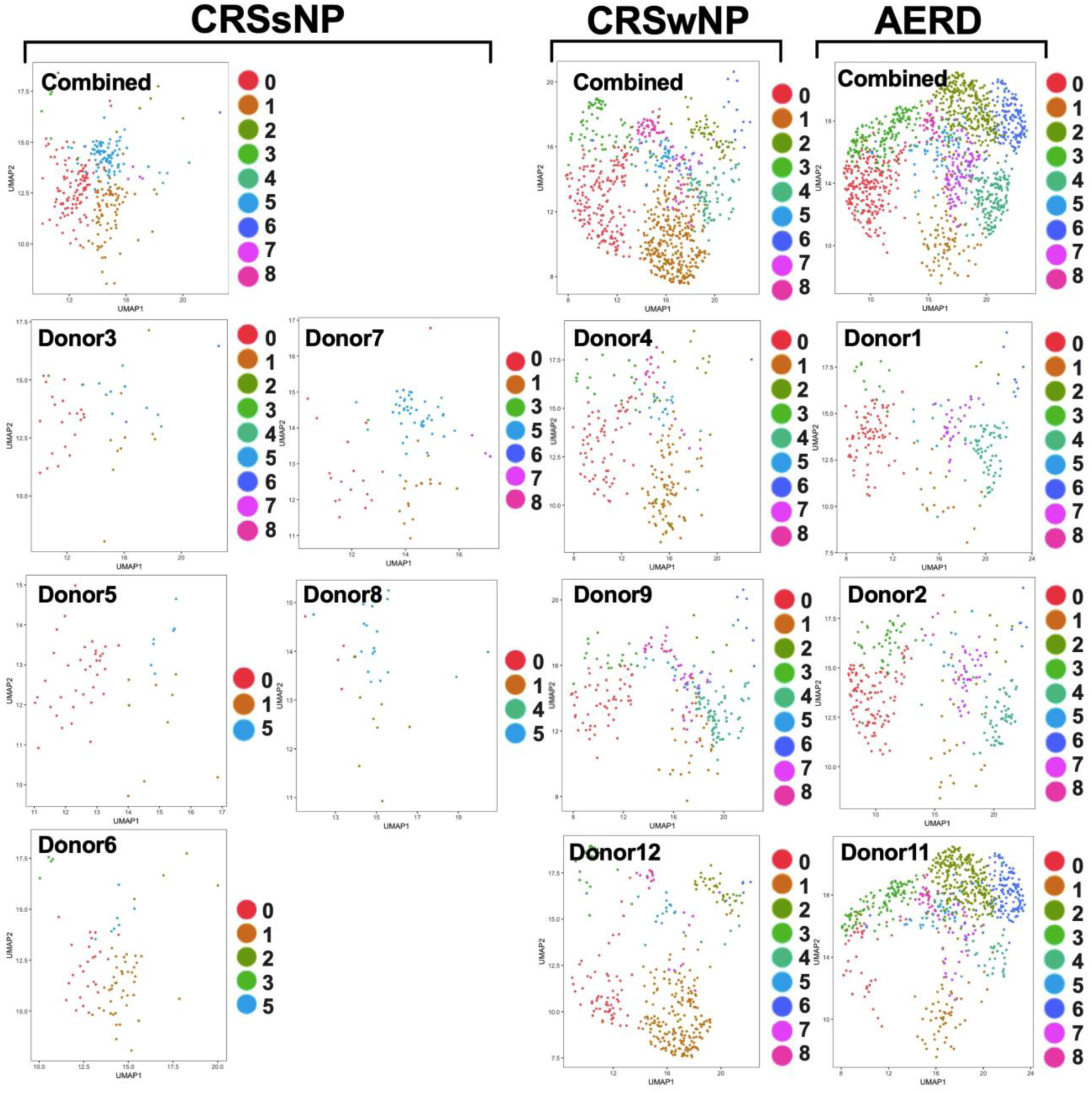
UMAP plots of 2,520 antibody-secreting cells colored by cluster as identified in **Fig. 3** and separated by disease (CRSsNP: 297 cells, CRSwNP: 835 cells, AERD: 1388 cells) and individual donor (Donor 1: 226 cells, Donor 2: 398 cells, Donor 3: 46 cells, Donor 4: 232 cells, Donor 5: 52 cells, Donor 6: 83 cells, Donor 7: 84 cells, Donor 8: 32 cells, Donor 9: 242 cells, Donor 11: 764 cells, Donor 12: 361 cells).

**Supplementary Figure 3.**
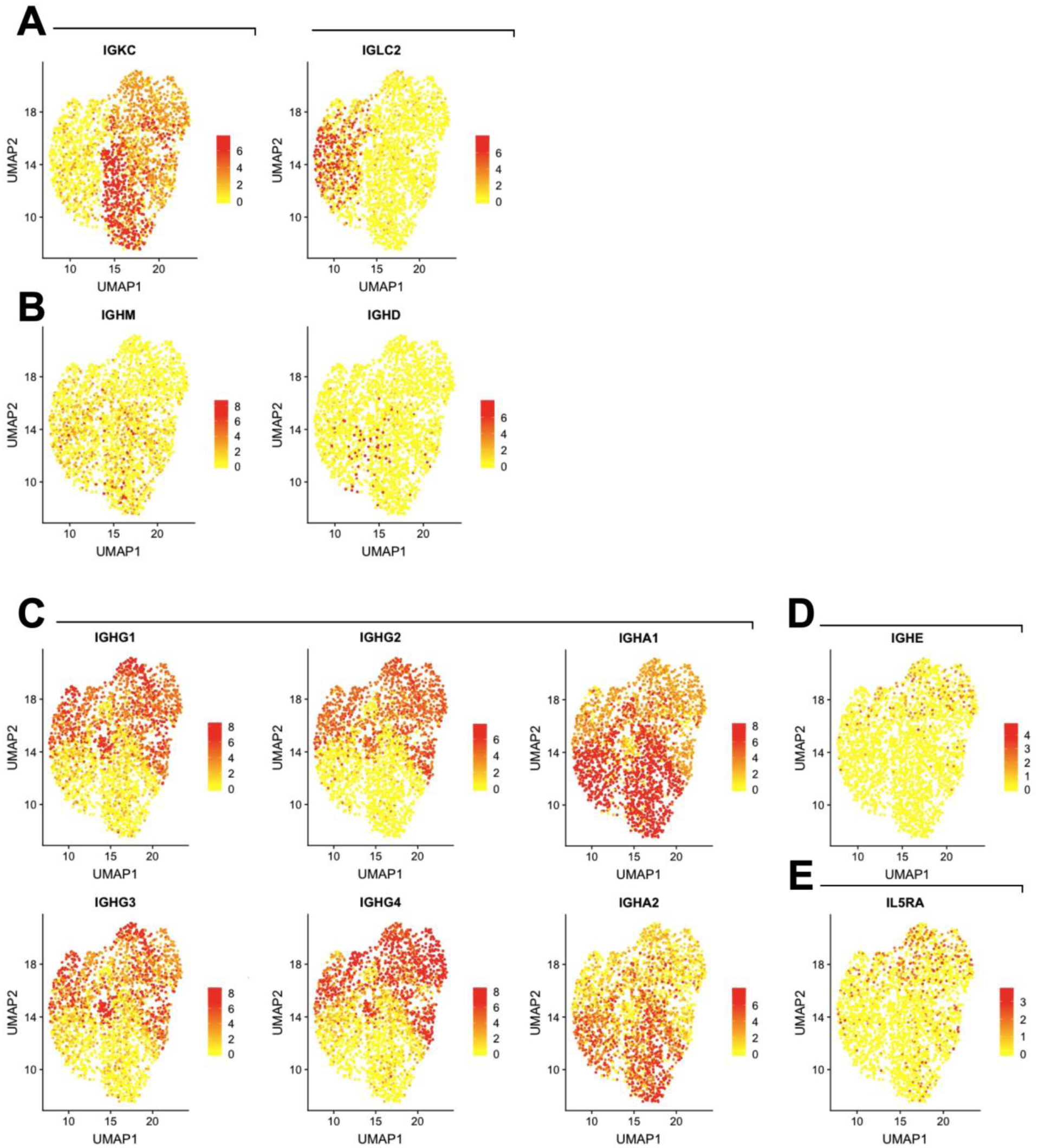
UMAP of 2,520 antibody-secreting cells, colored by expression of (**A**) Light chain constant regions (*IGKC*, *IGLC2*), (**B**) Non-class-switched heavy chain constant regions (*IGHM*, *IGHD*), (**C**) class-switched IgG and IgA isotype heavy chain regions (*IGHG1*, *IGHG2*, *IGHG3*, *IGHG4*, *IGHA1*, *IGHA2*), (**D**) the immunoglobulin E heavy chain (*IGHE*), and (**E**) the IL-5 receptor alpha chain (*IL5RA*). Scale indicates log-normalized expression.

**Supplementary Figure 4.**
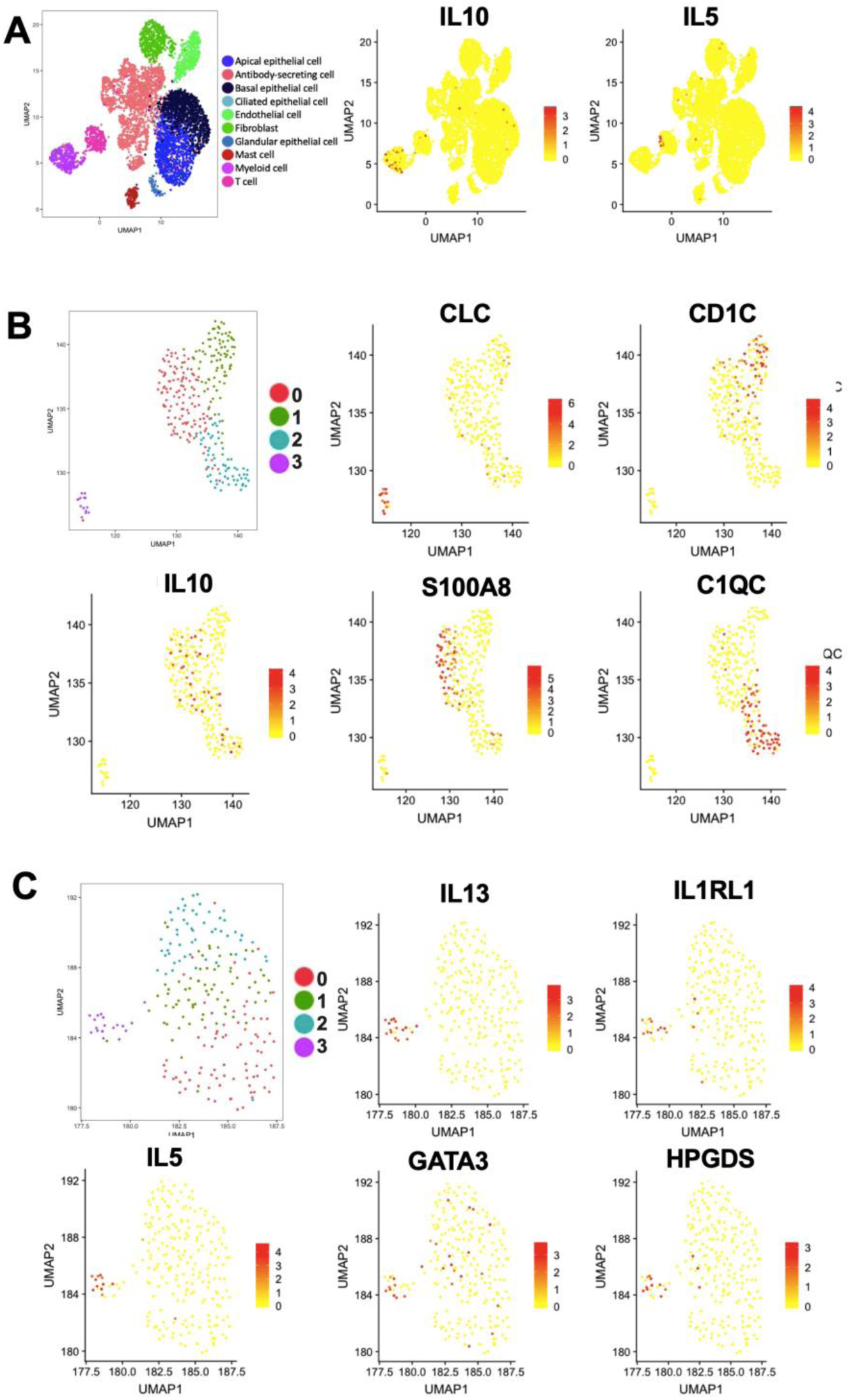
UMAP plot of 4,276 nasal polyp cells from subjects with AERD (n=3 samples), colored by cell type (left) and IL5 and IL10 expression (right) (**B**) UMAP plot of 282 myeloid lineage derived from (A), clustered with accompanying plots to show *IL10* expression across sub-populations including *S100A8*-expressing inflammatory DC3 (Cluster 0) and *C1QC*-expressing macrophages (cluster 2), but not including *CLC-* expressing eosinophils (cluster 3) or *CD1C*-expressing DC2 (cluster1). (**C**) UMAP plot of 224 T lymphocytes derived from (A), clustered with accompanying plots to show *IL5* expression within cluster 3, which additionally expressed *IL13*, *IL1RL1*, *GATA3* and *HPGDS*, suggesting a Th2A phenotype. Scale indicates log-normalized expression.

**Supplemental Table E1.**
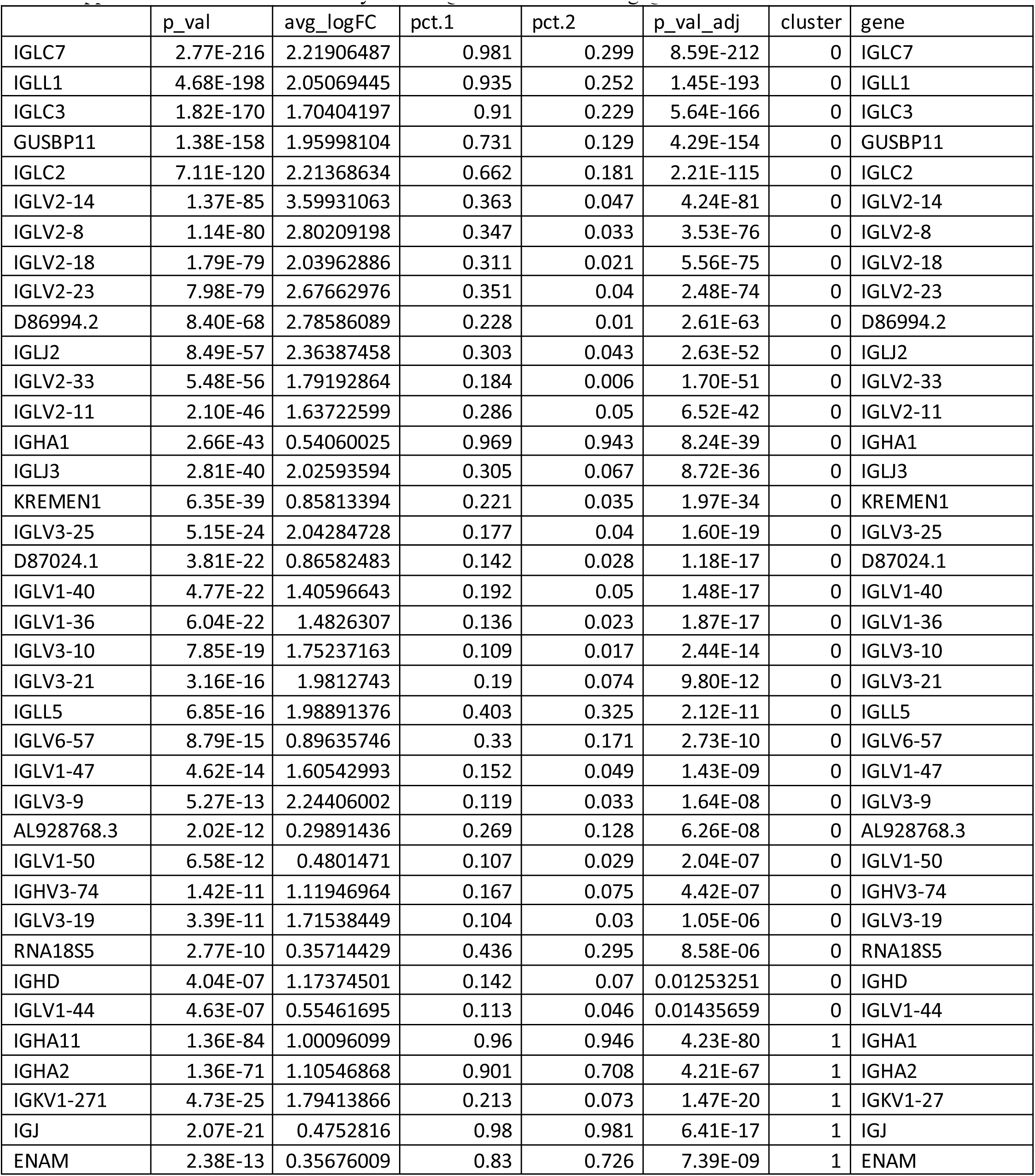
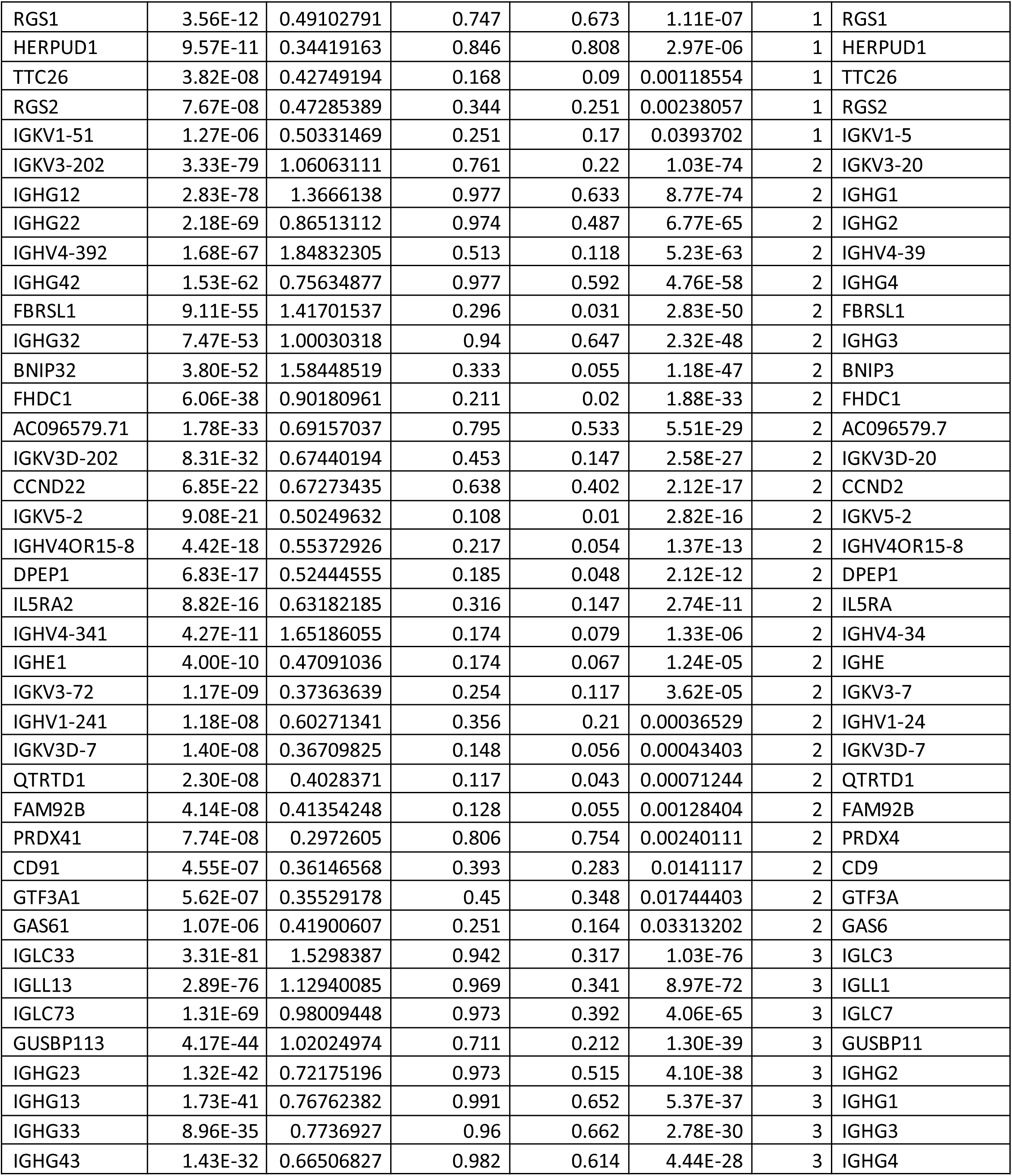
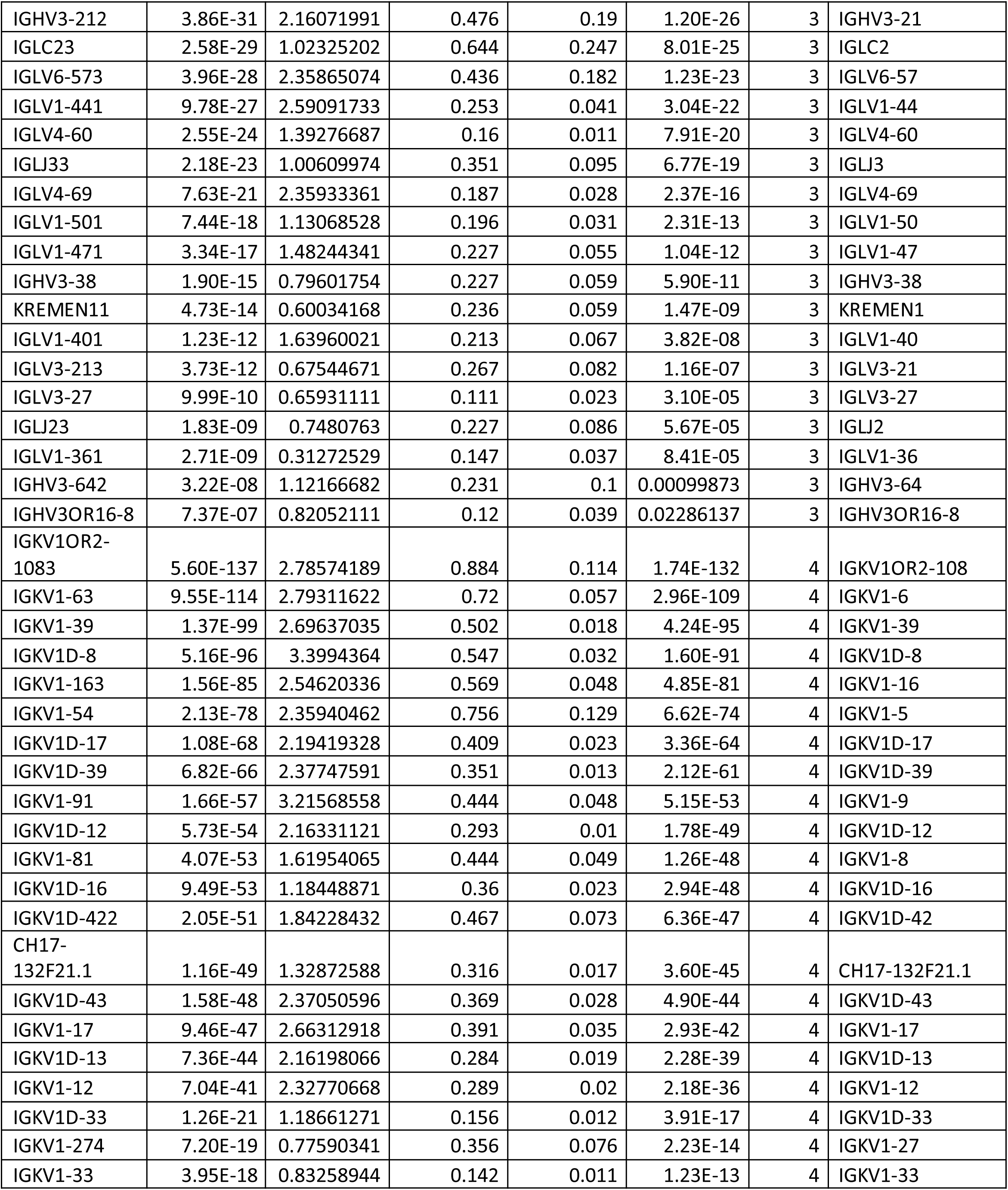
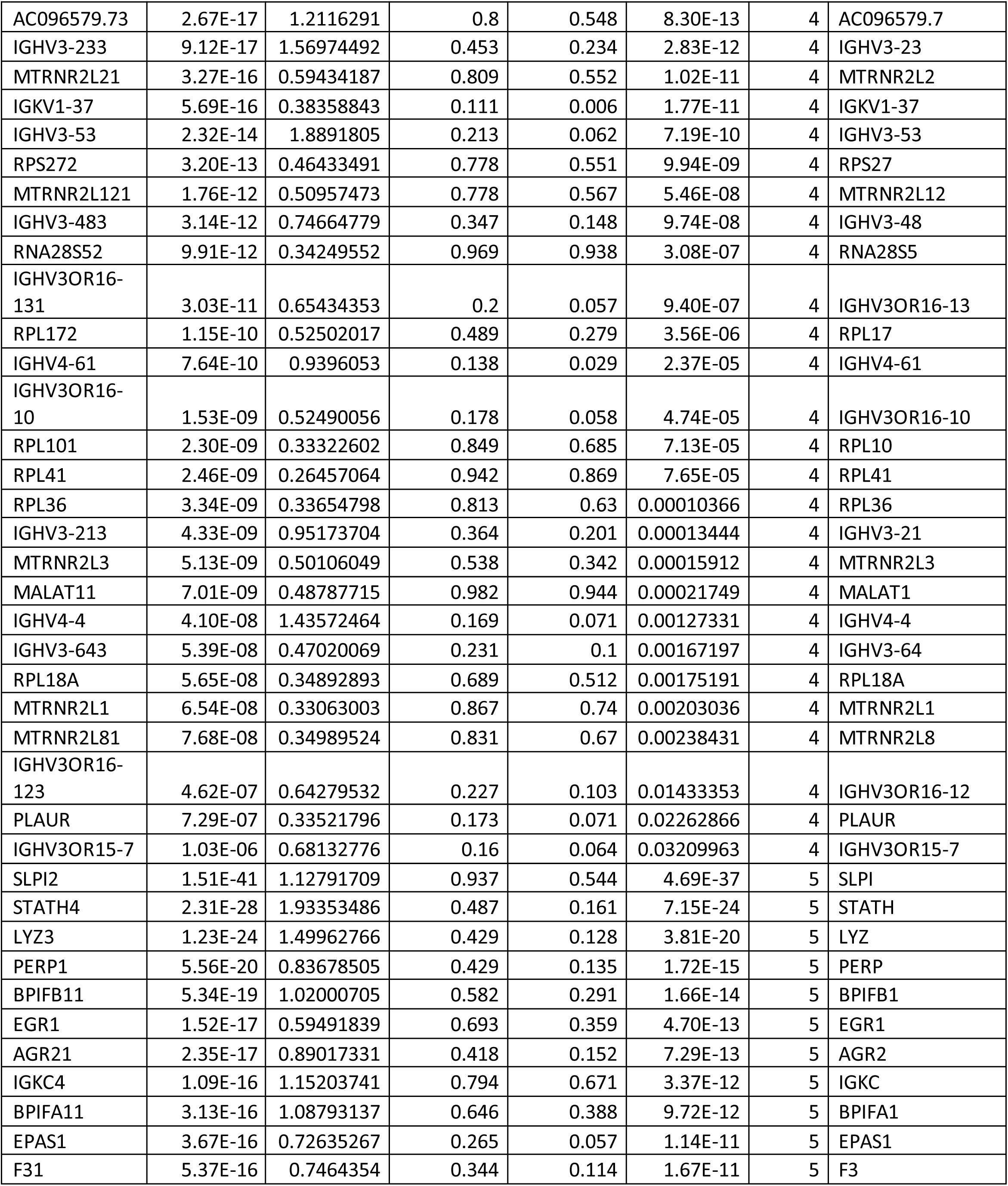
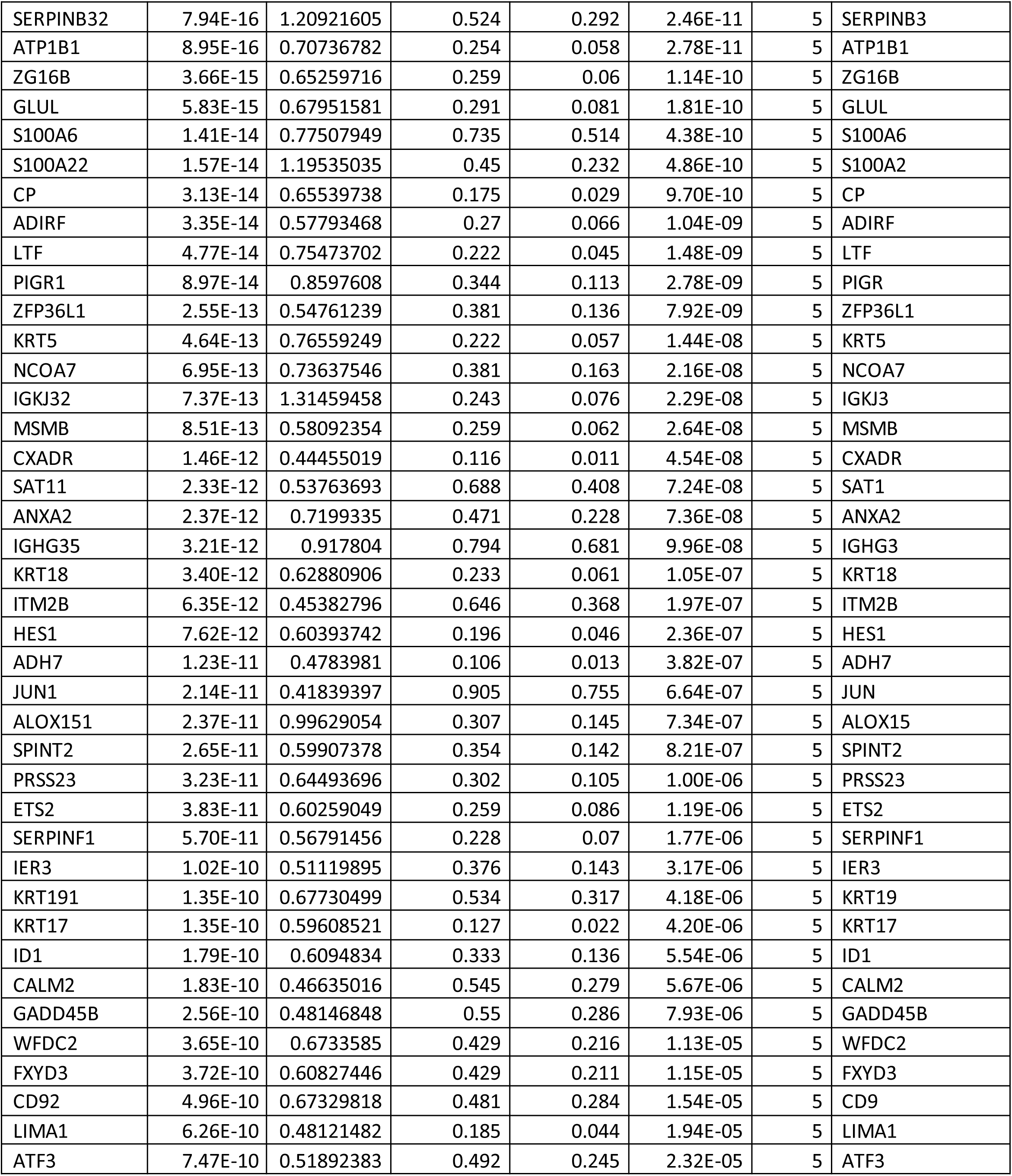
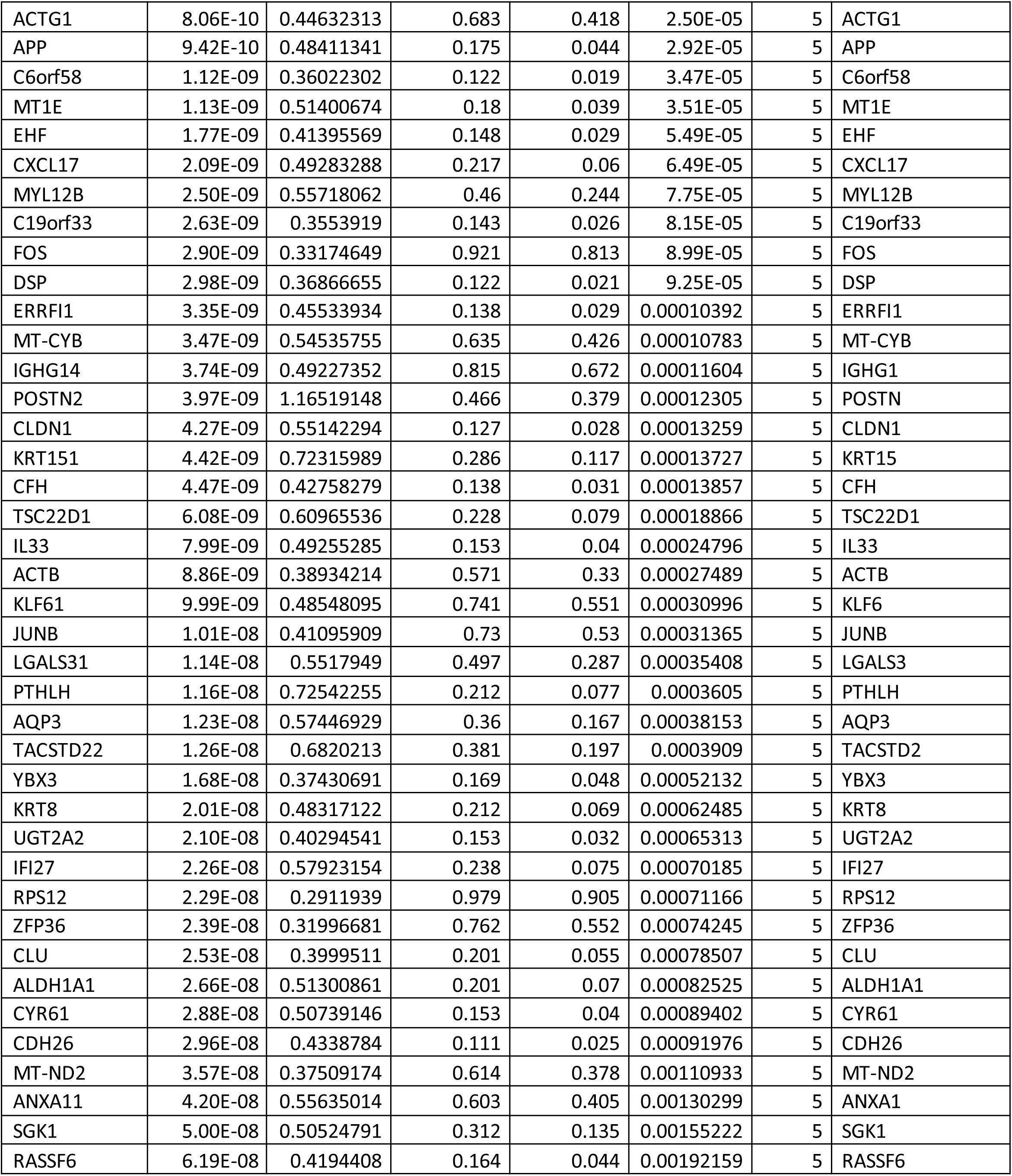
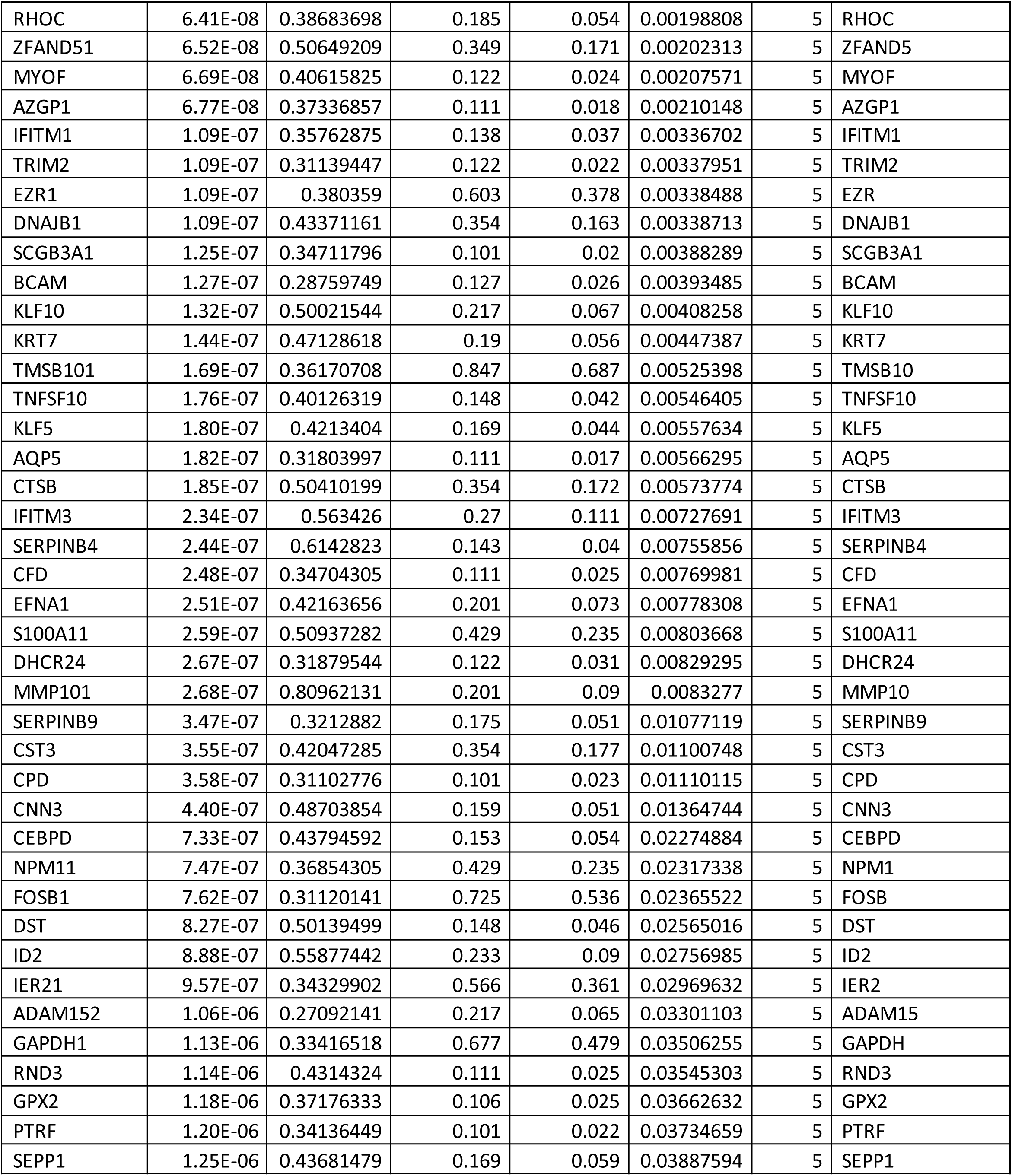
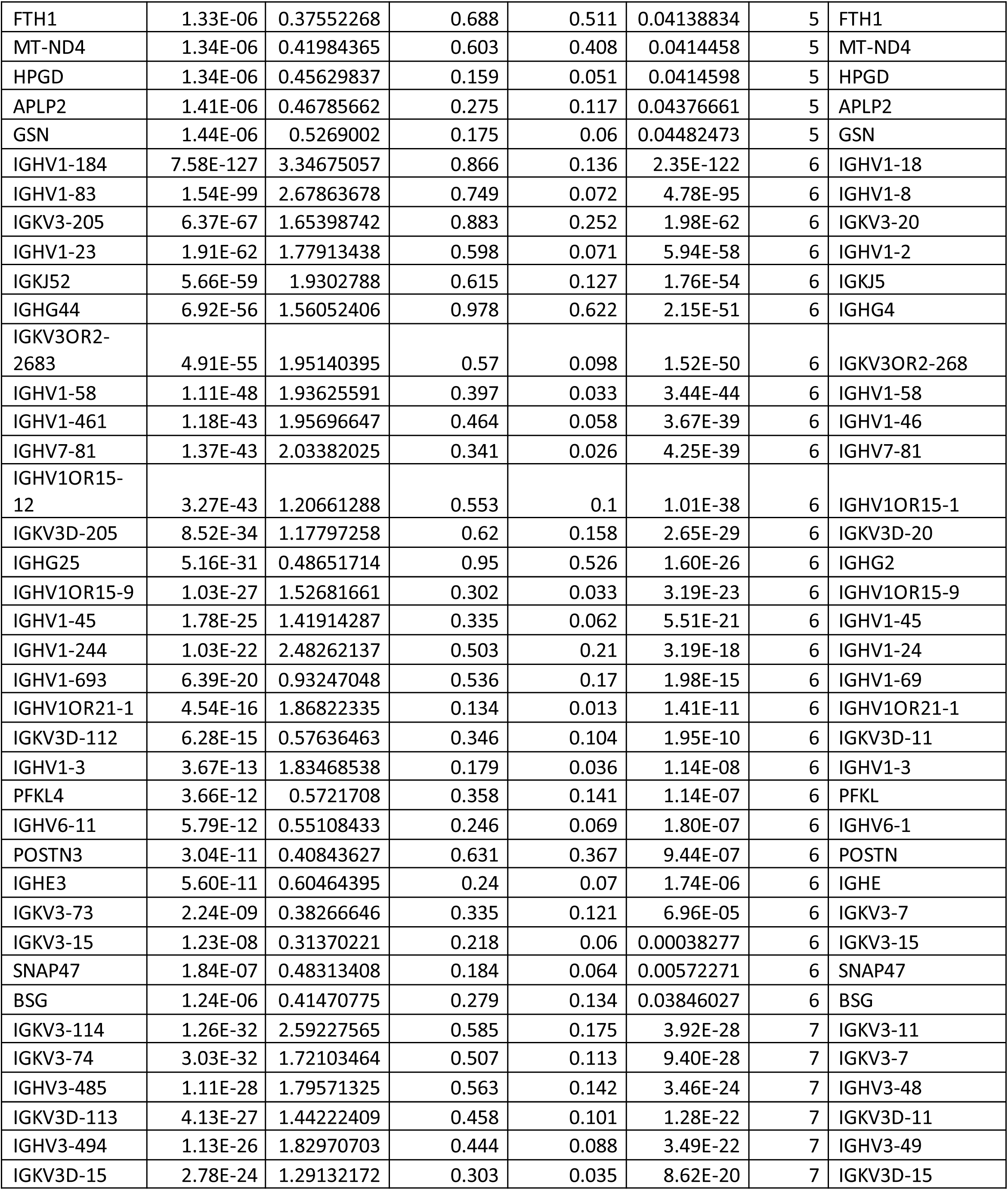
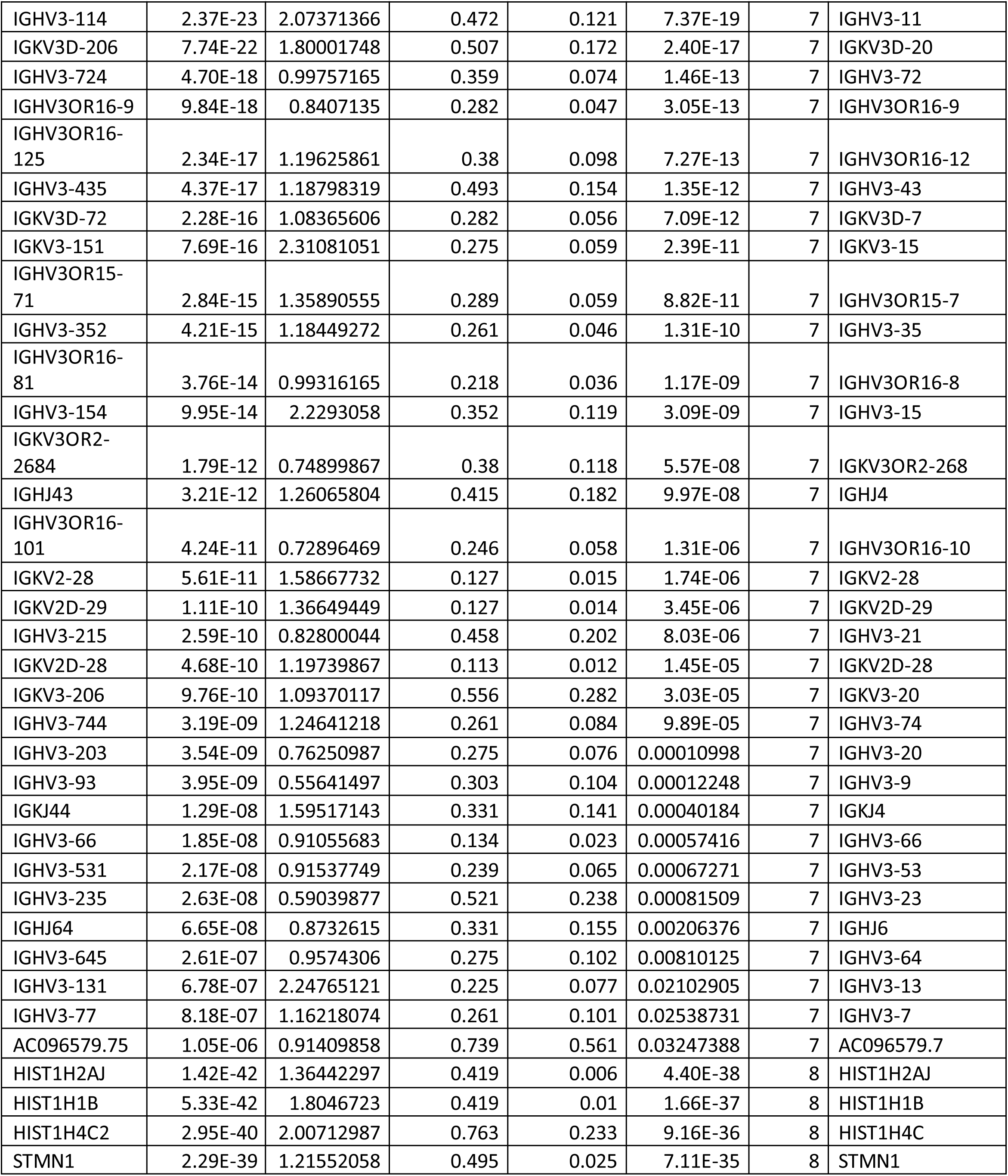
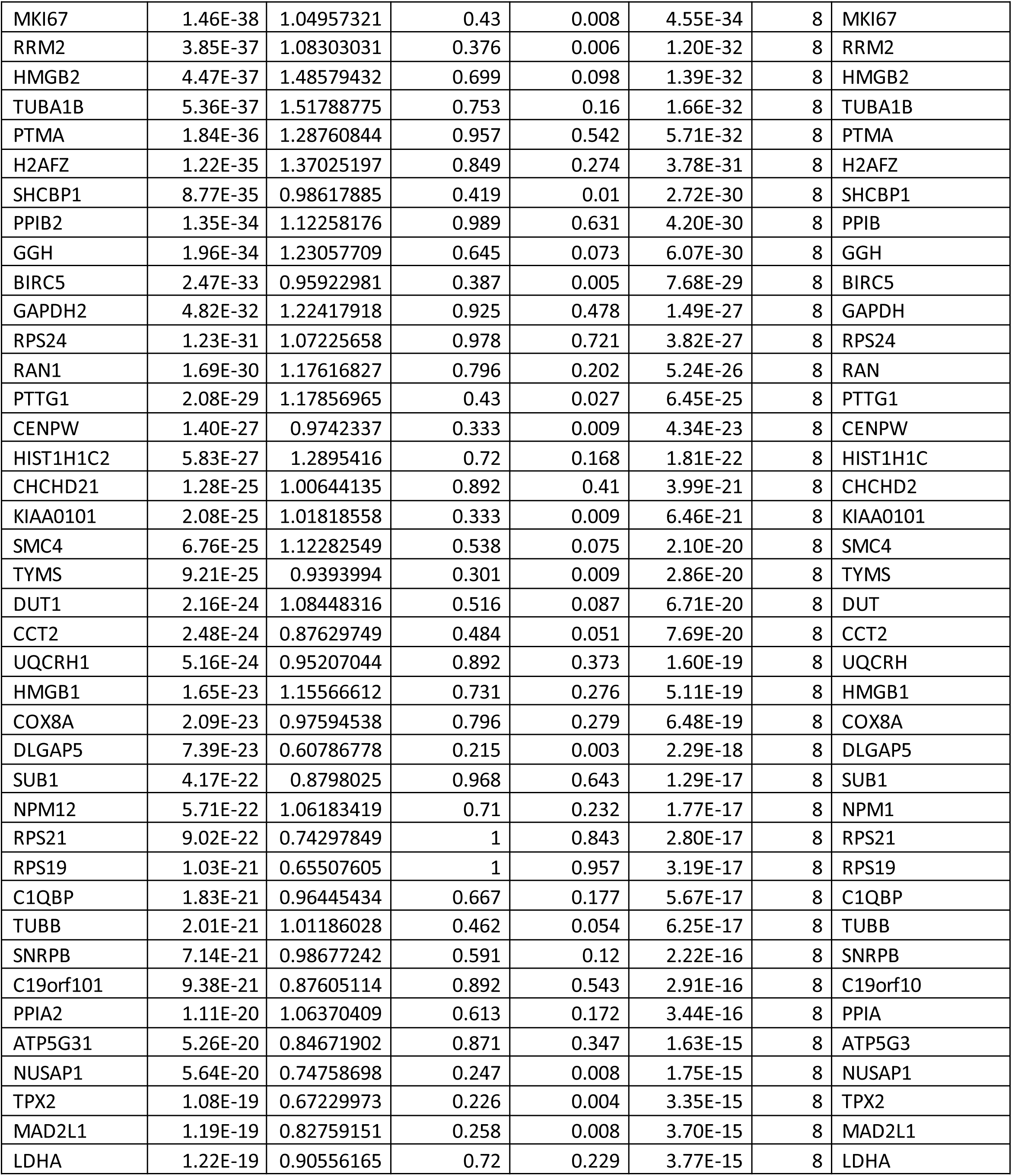
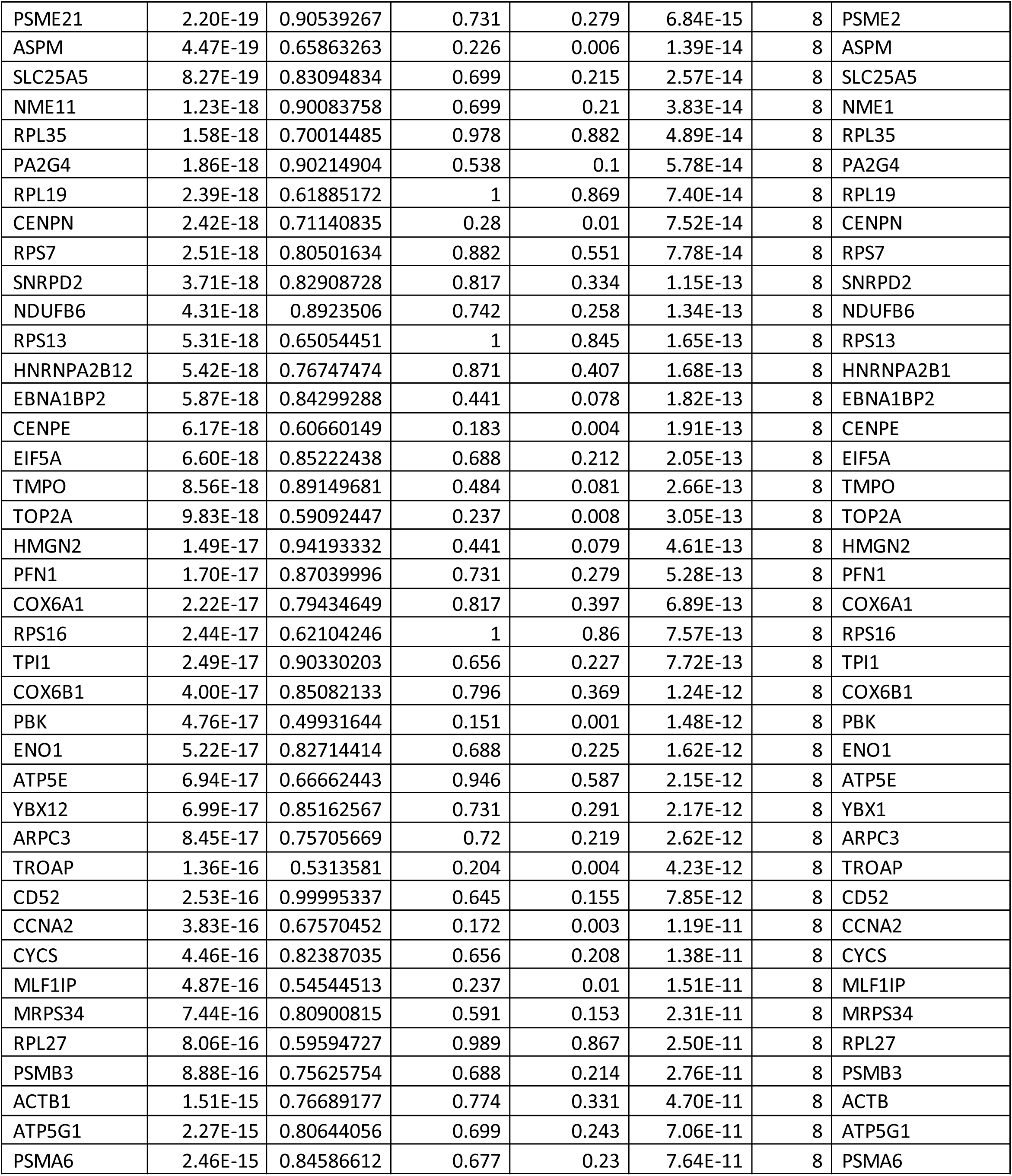
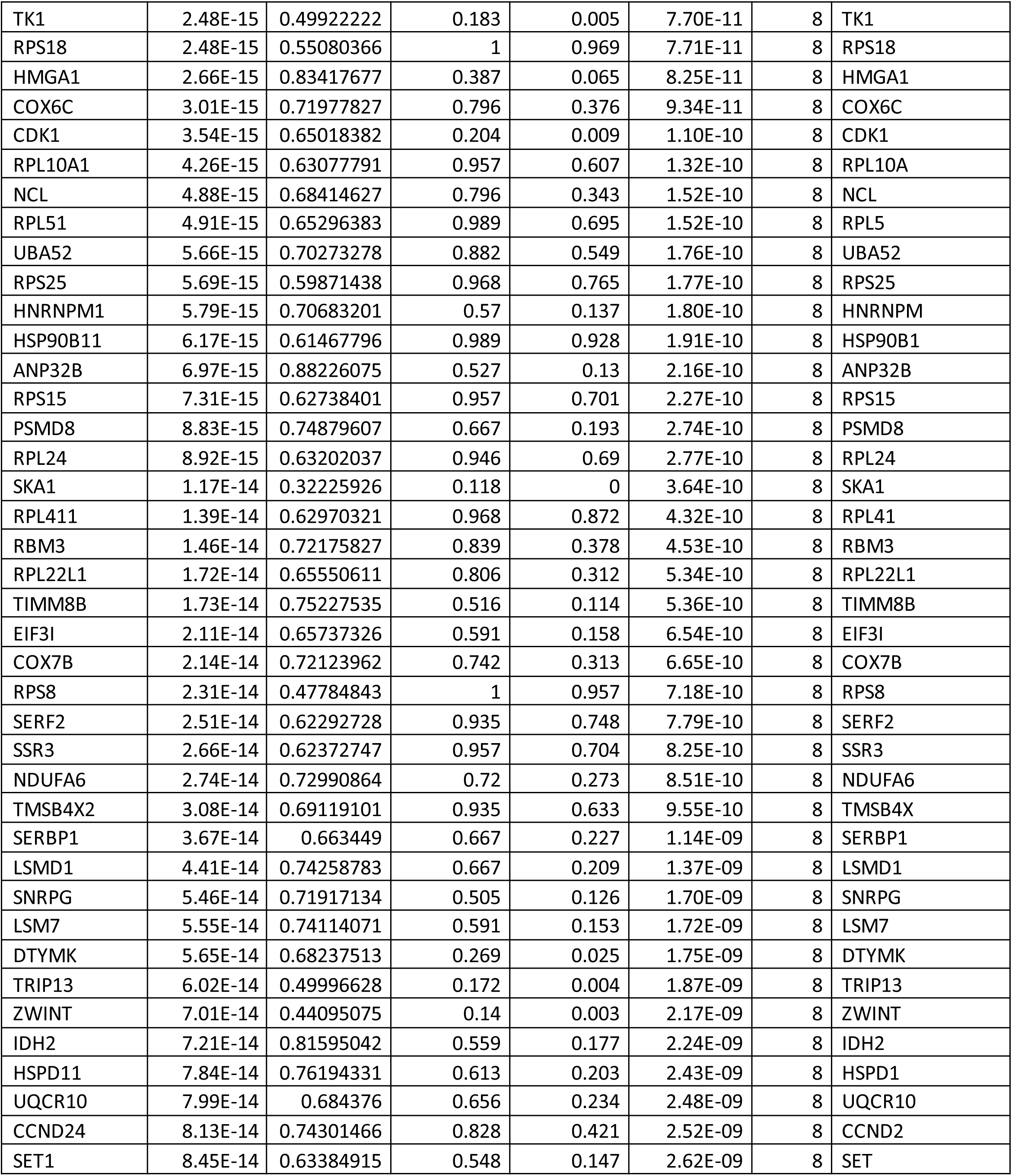
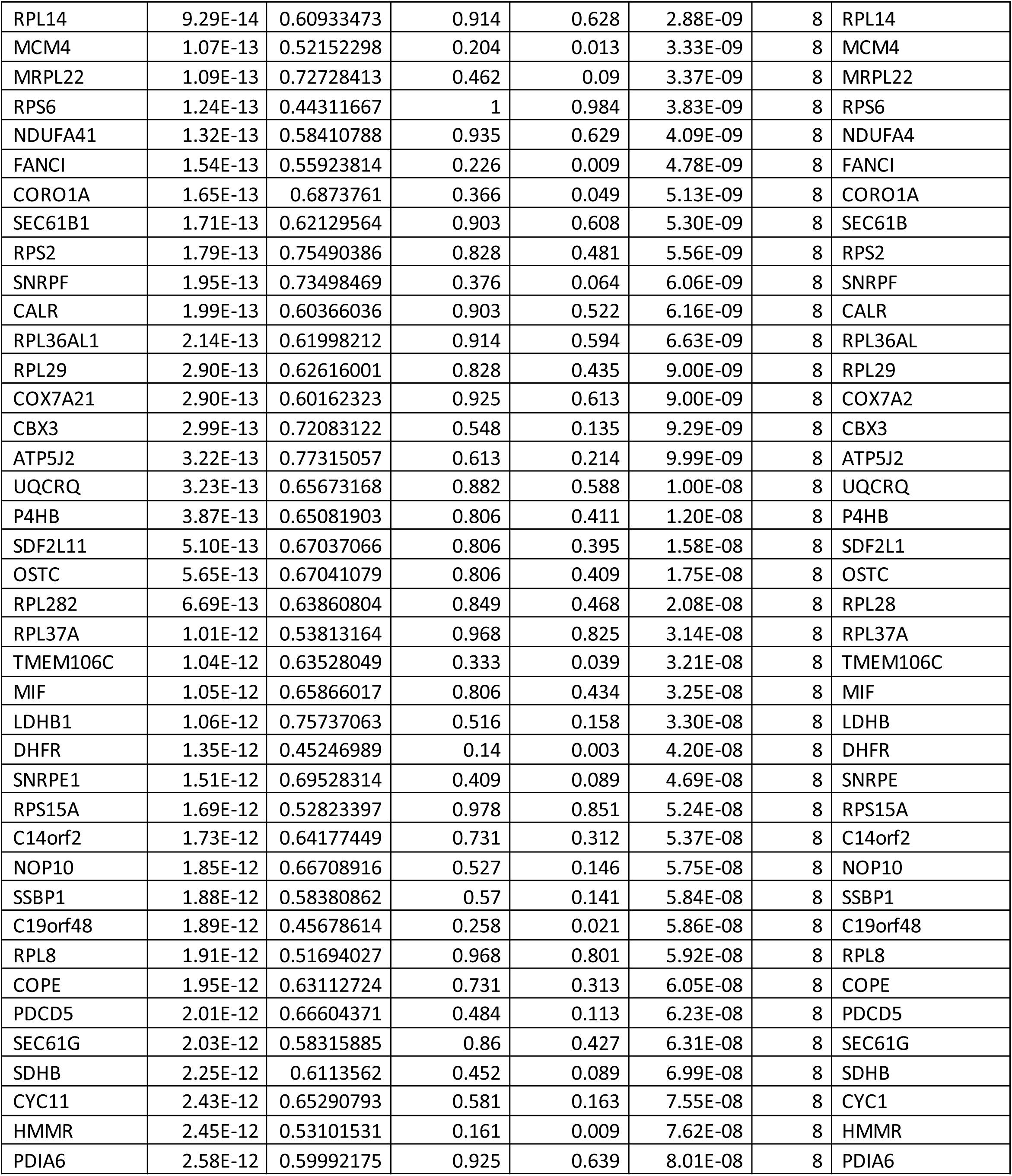
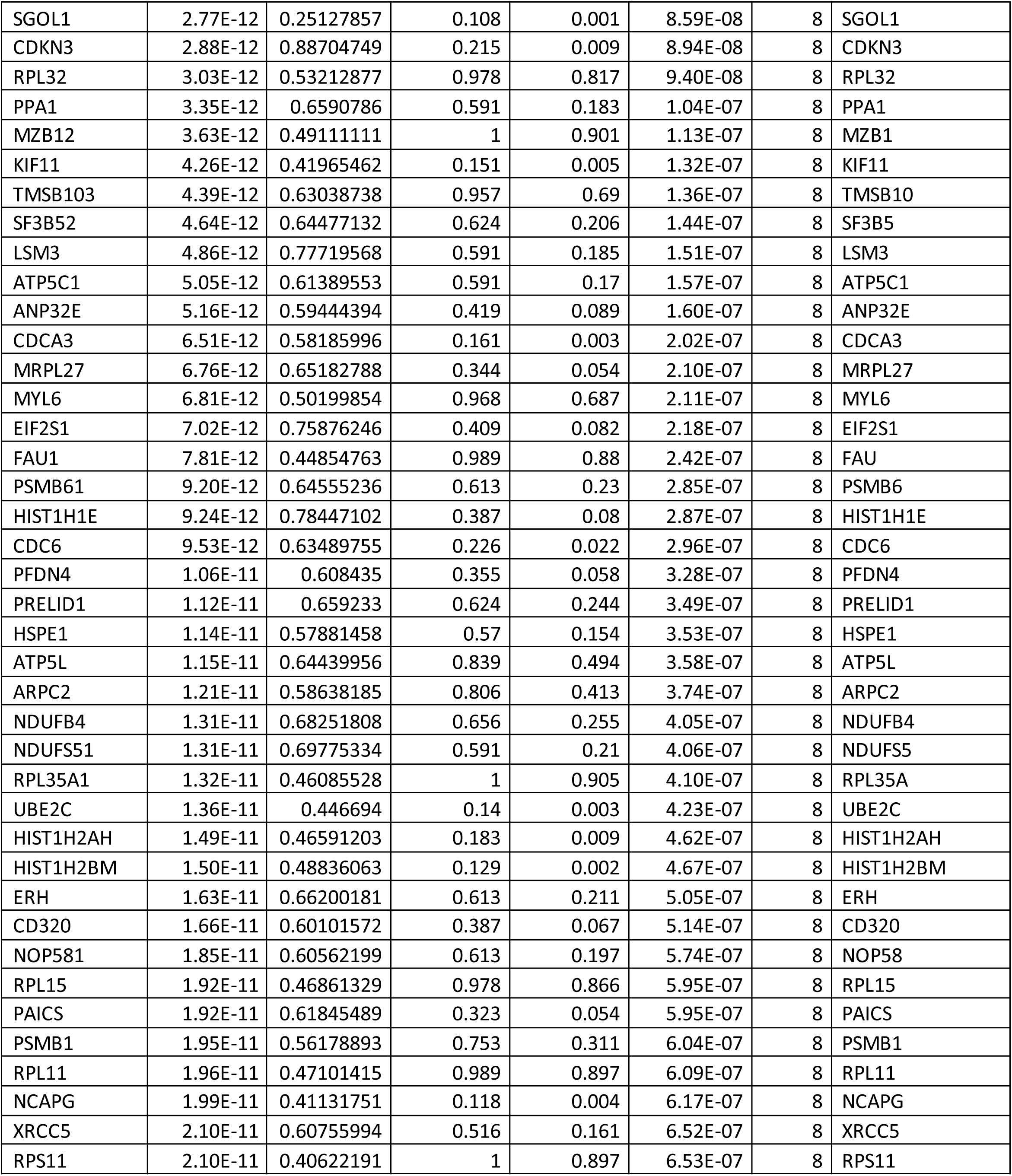
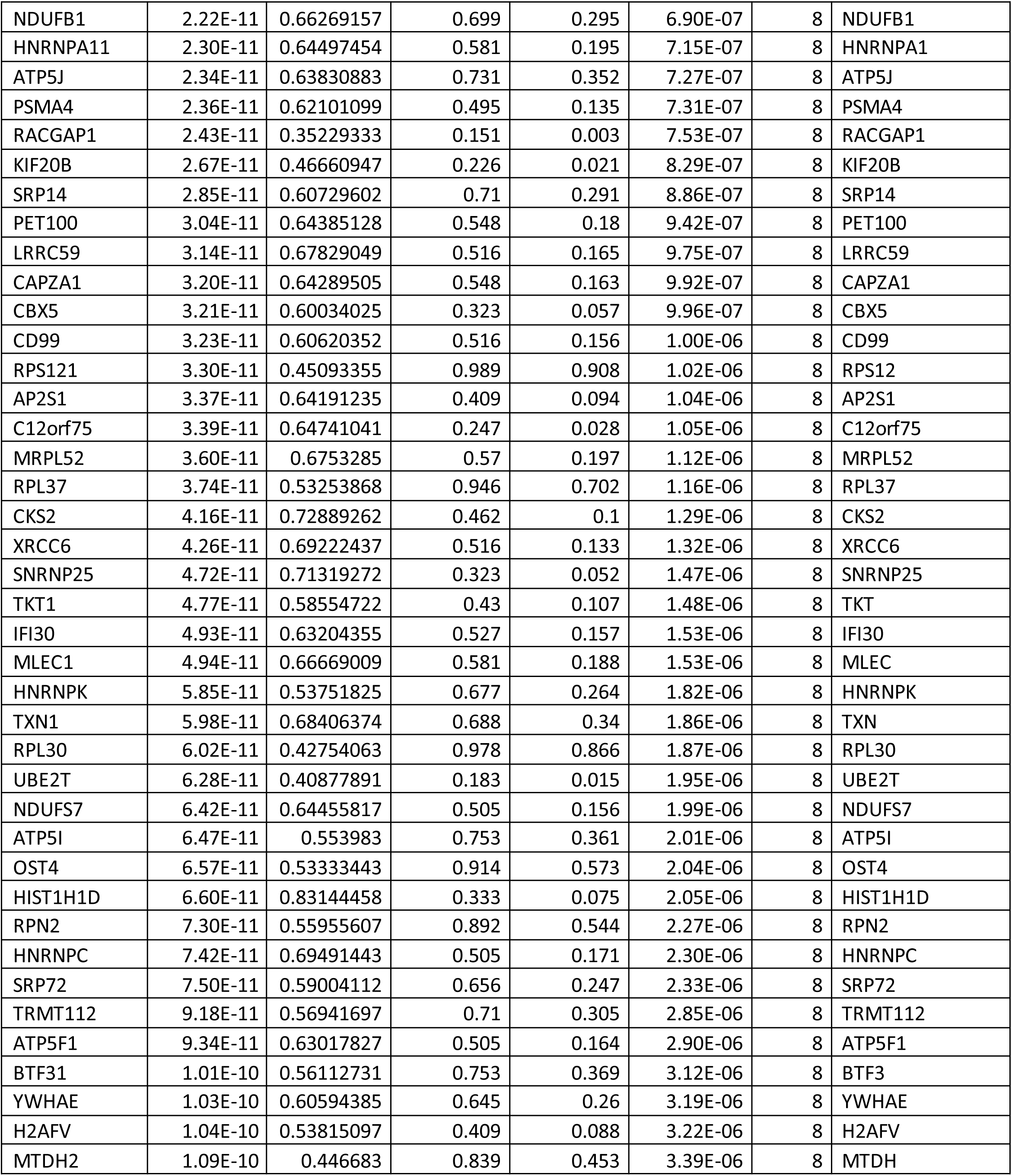
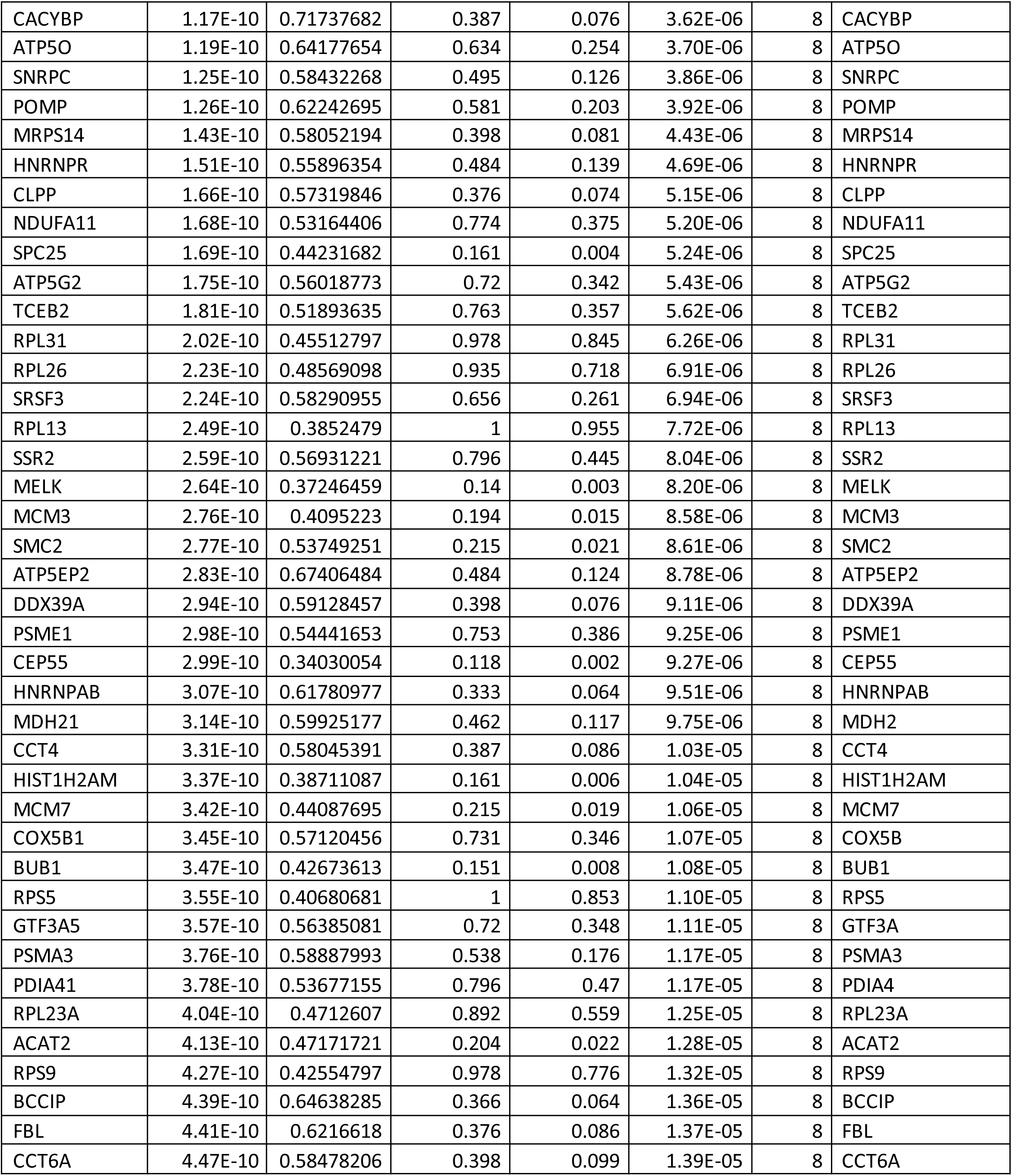
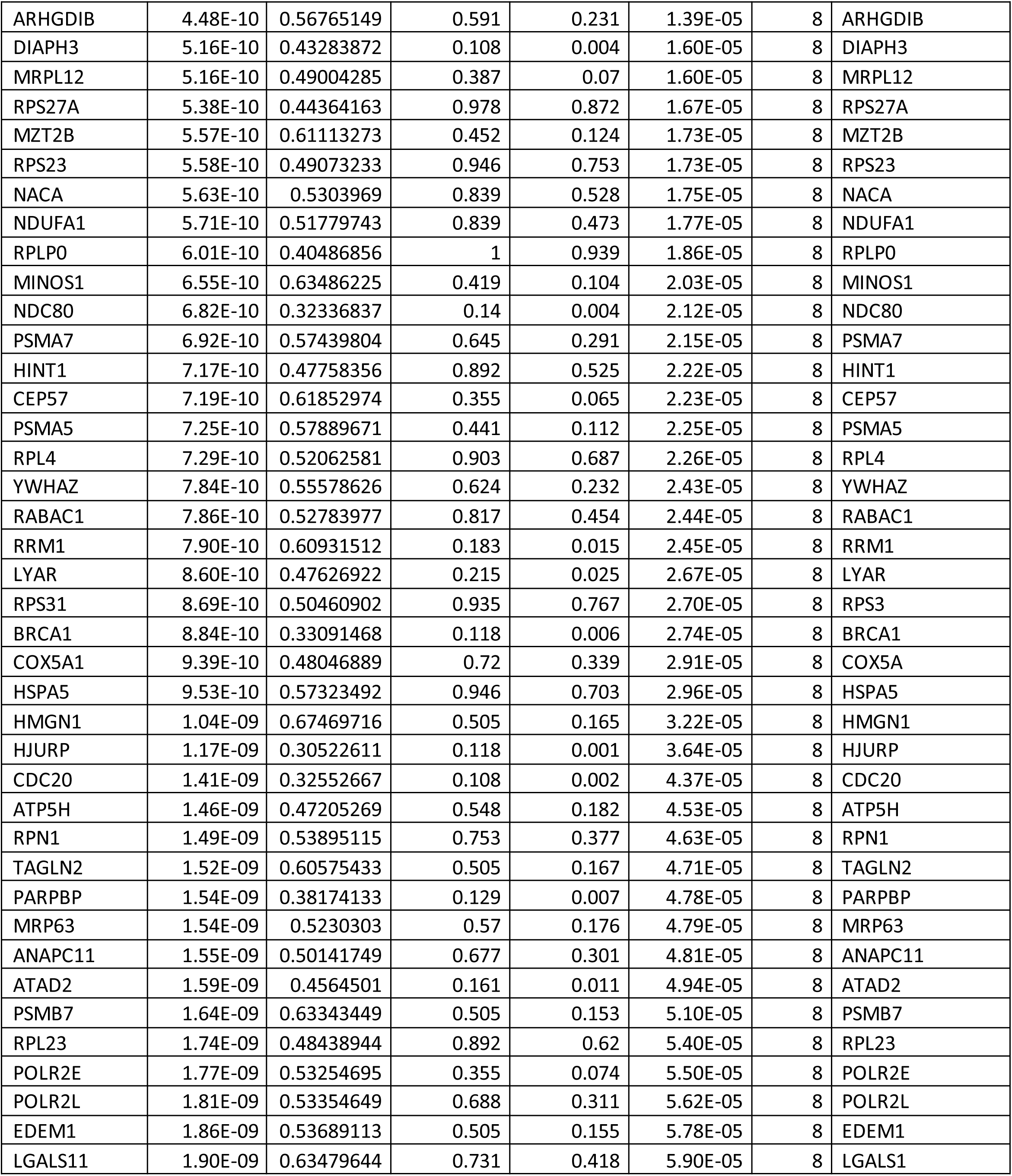
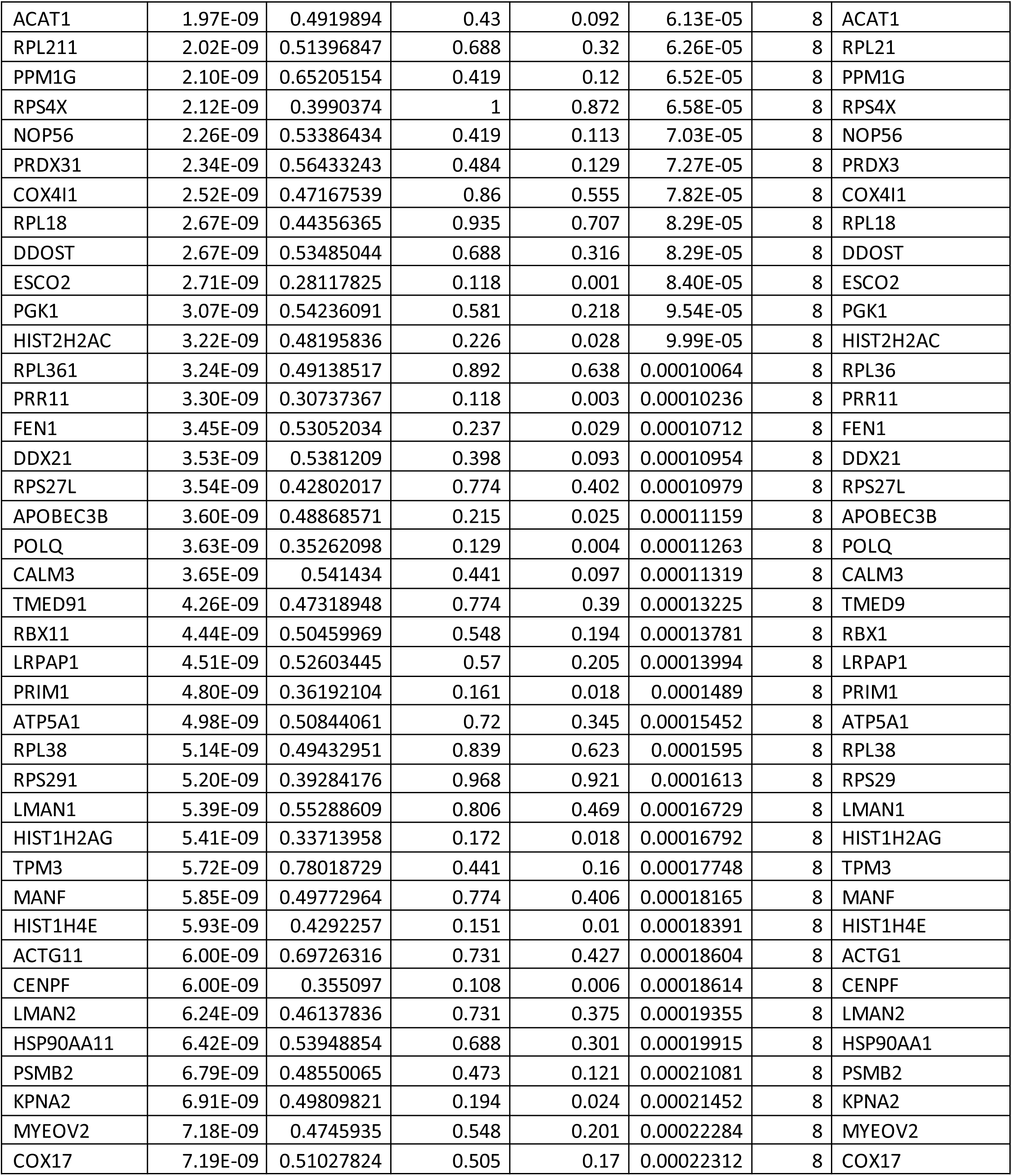
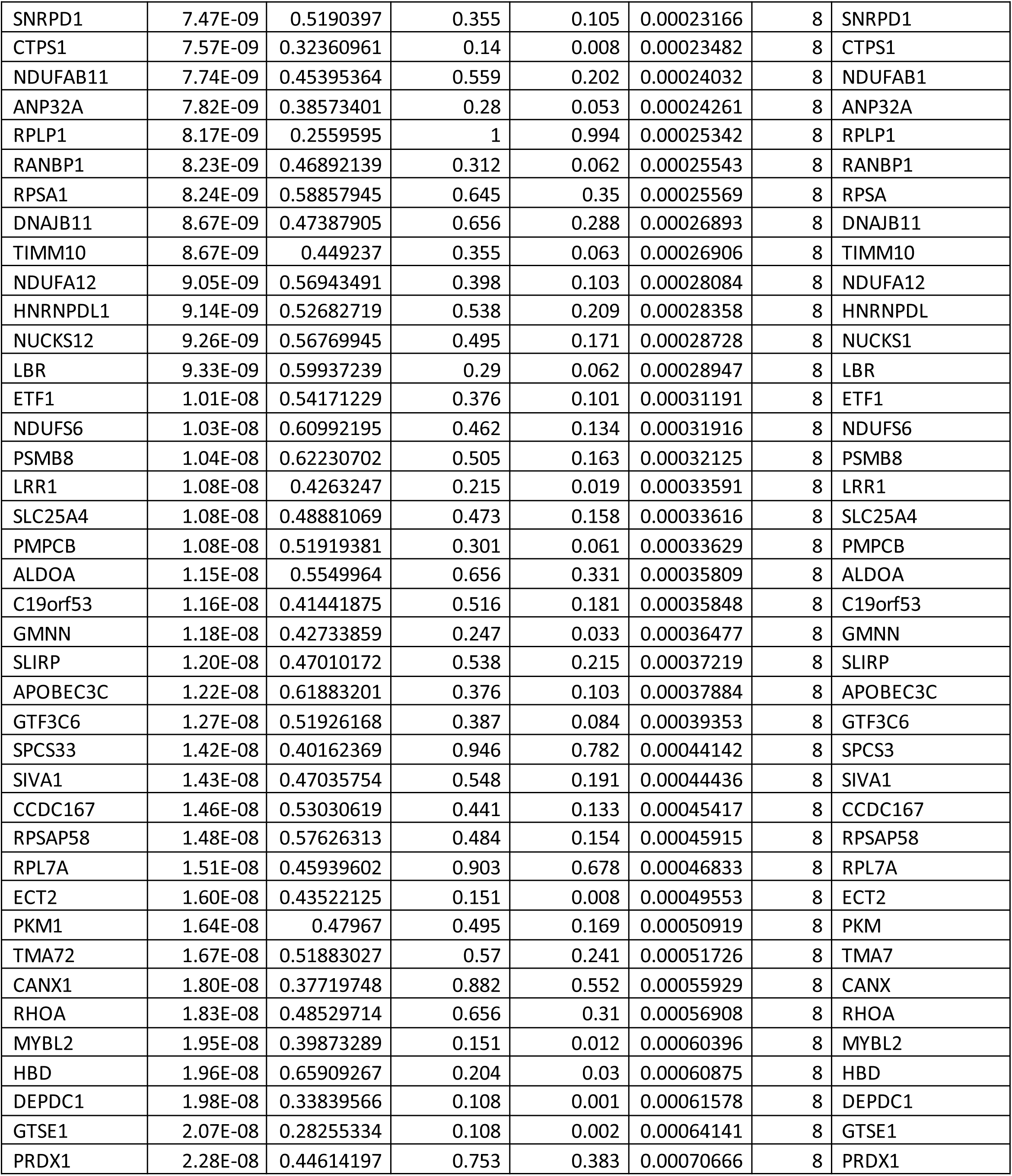
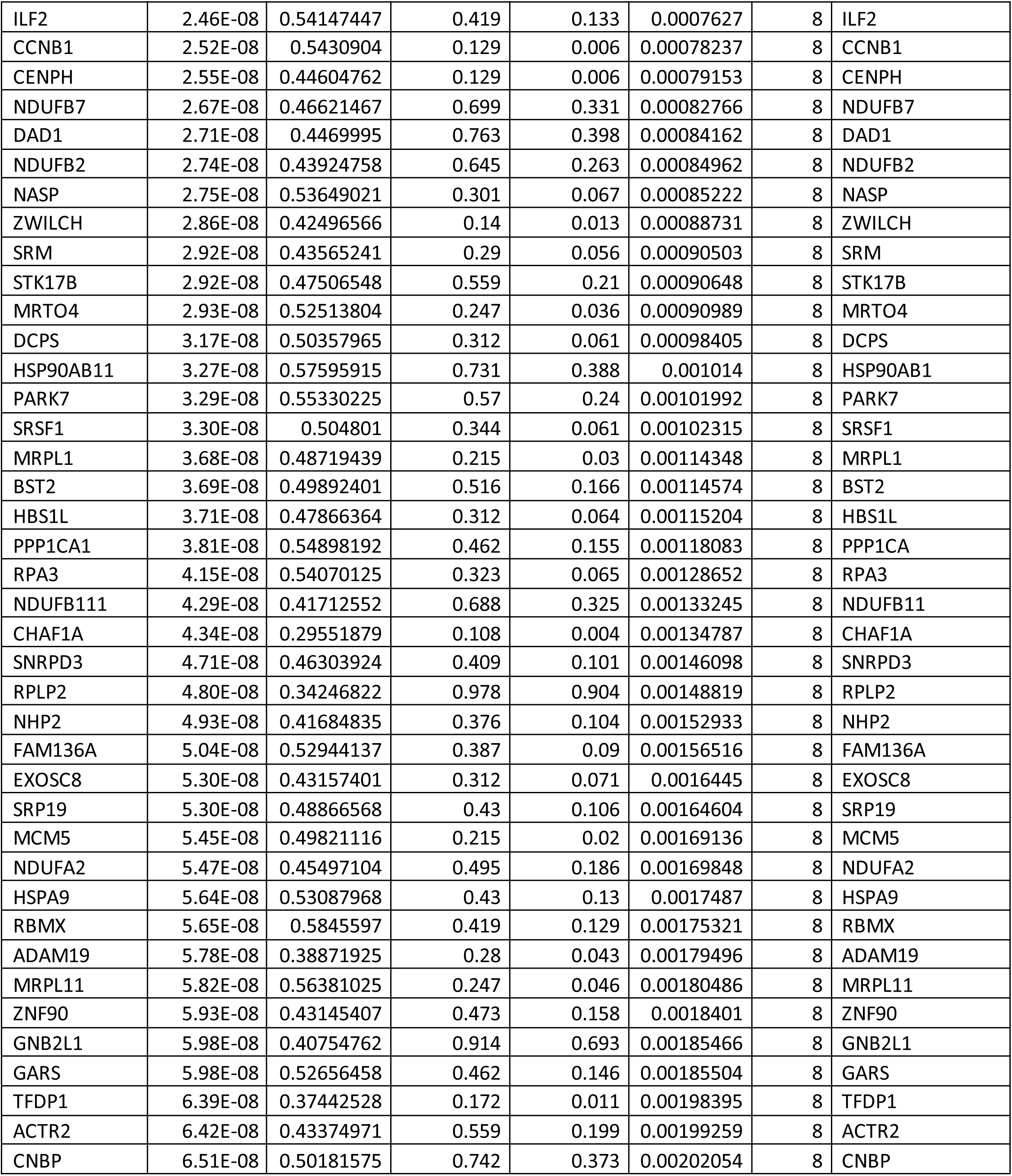
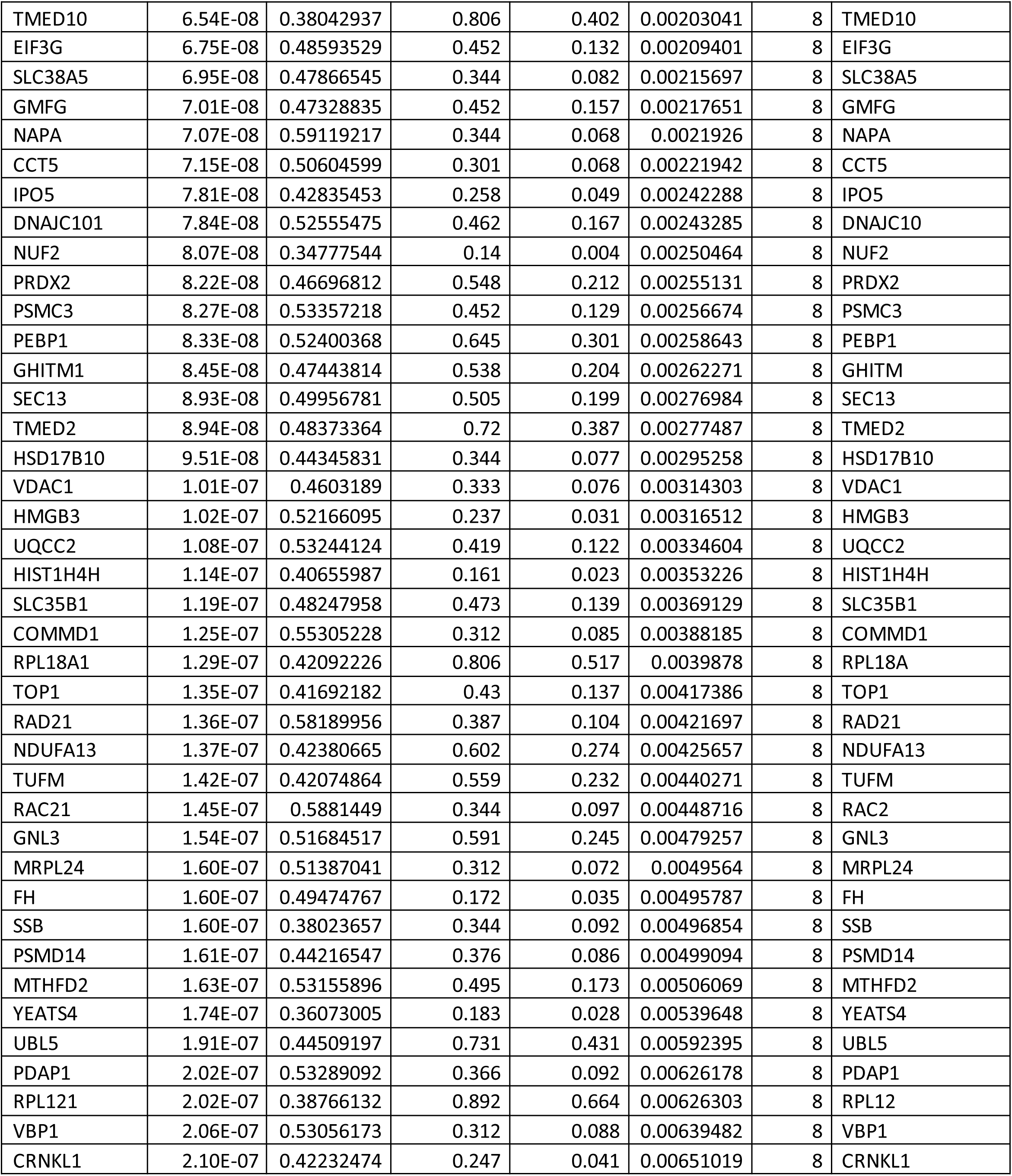
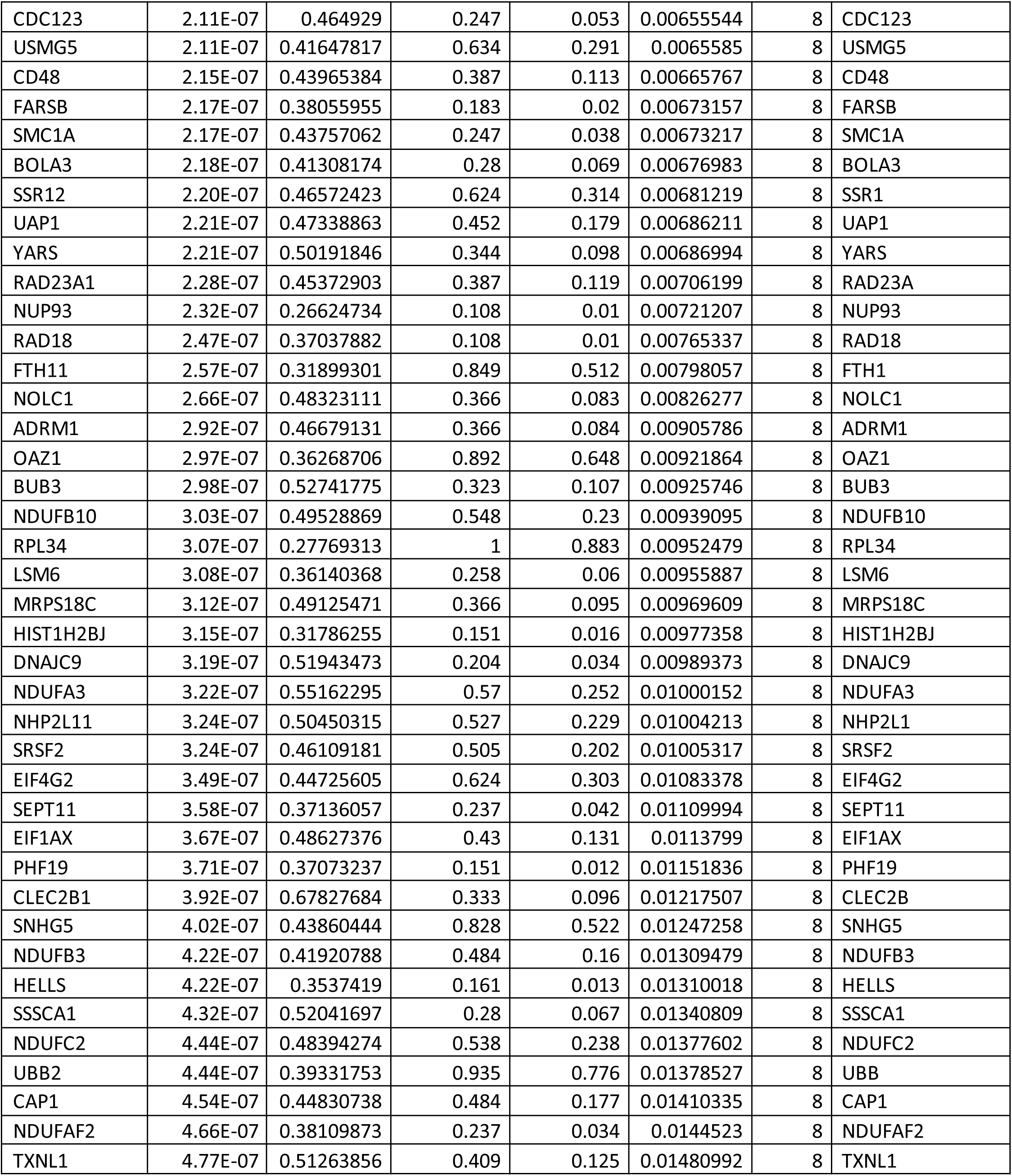
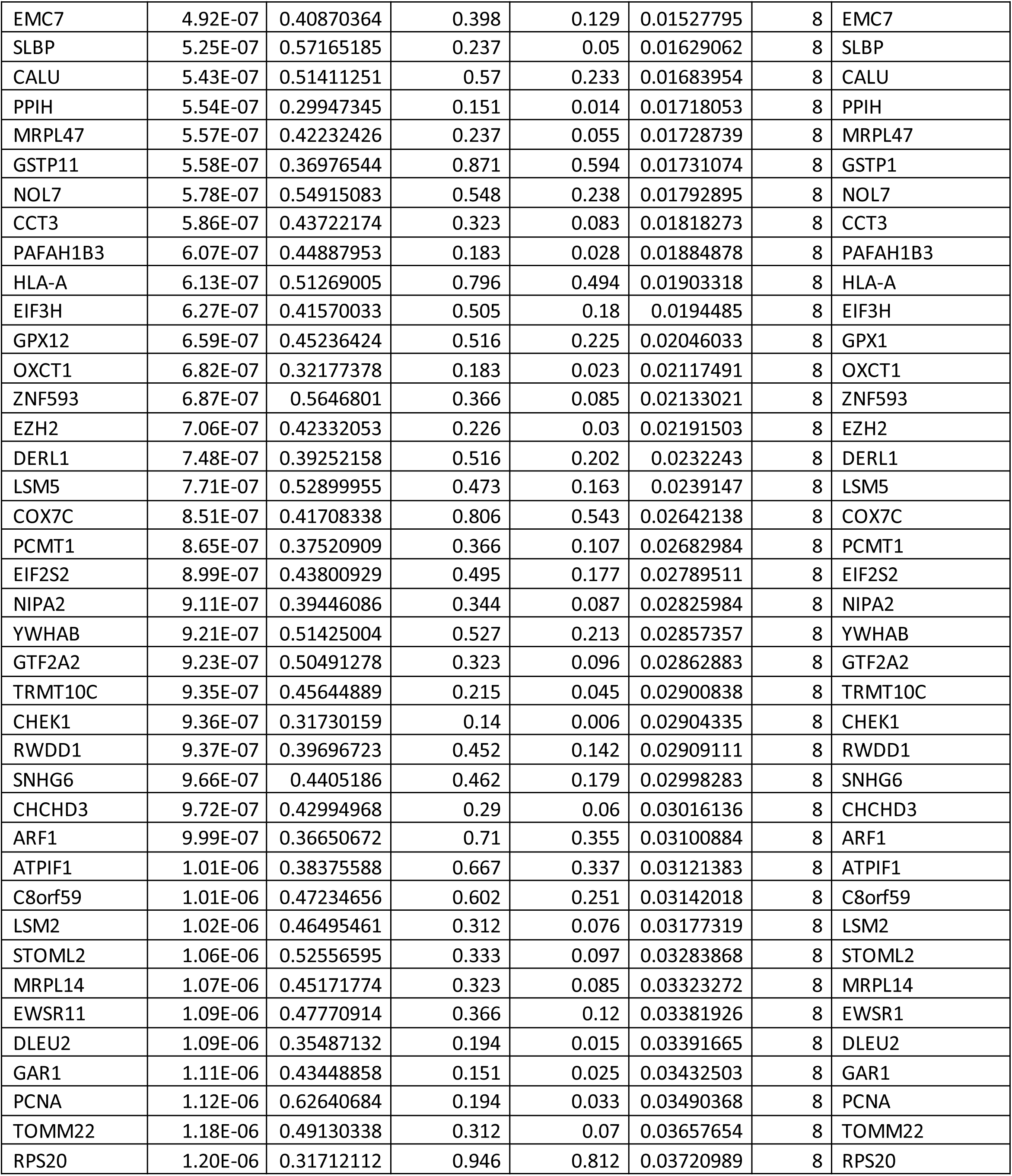
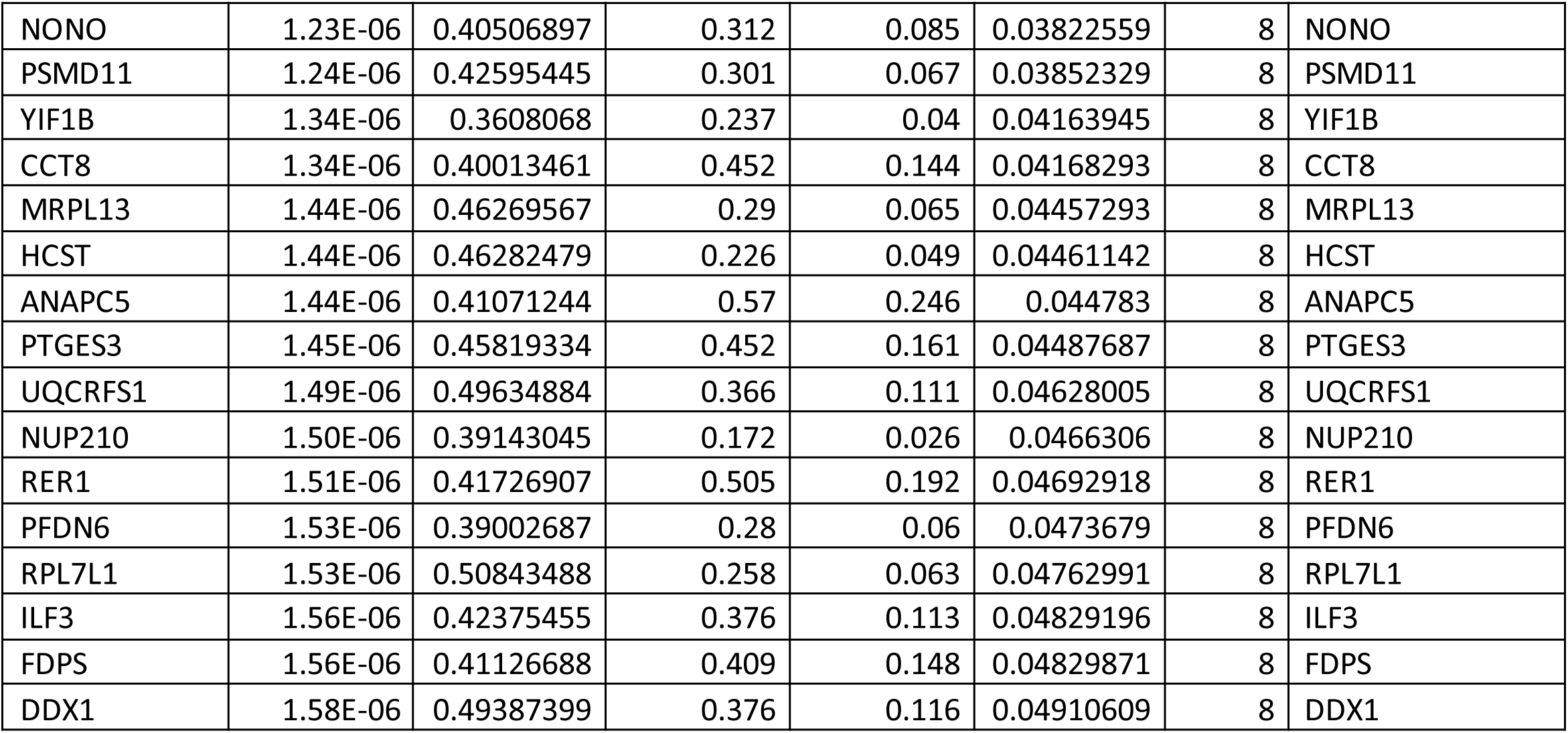
Antibody-secreting cell cluster defining genes.

**Supplemental Table E2.**
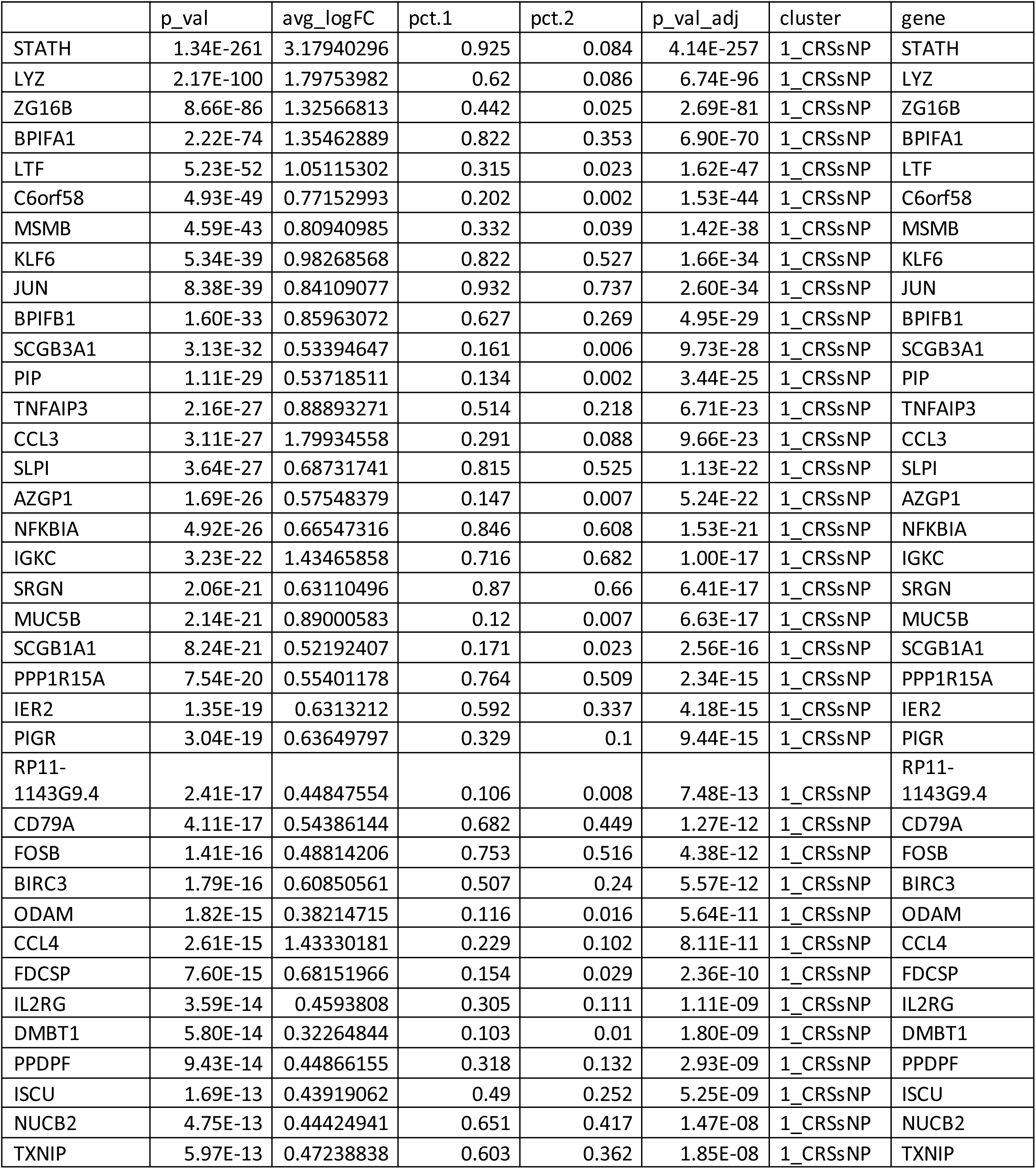
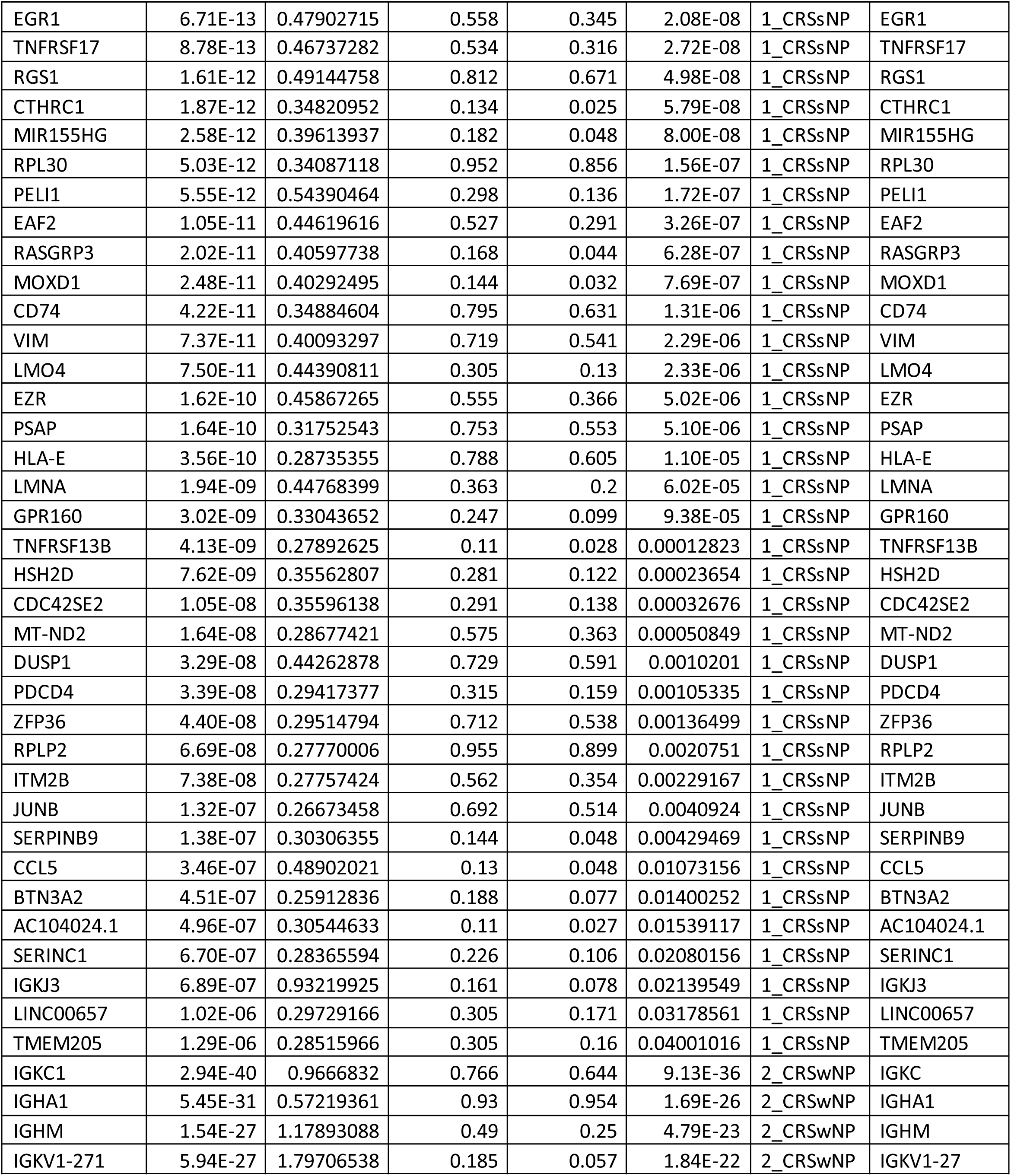
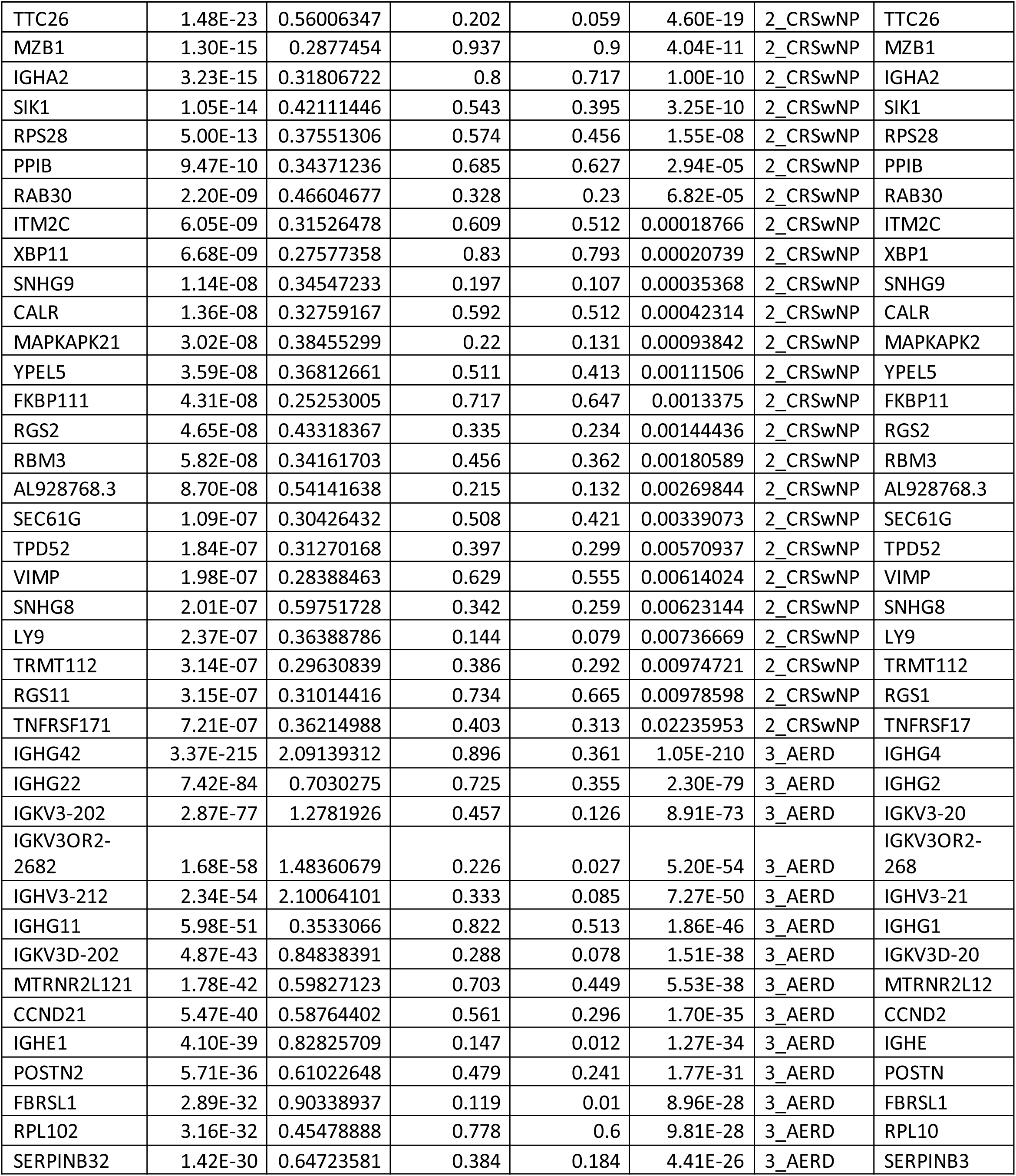
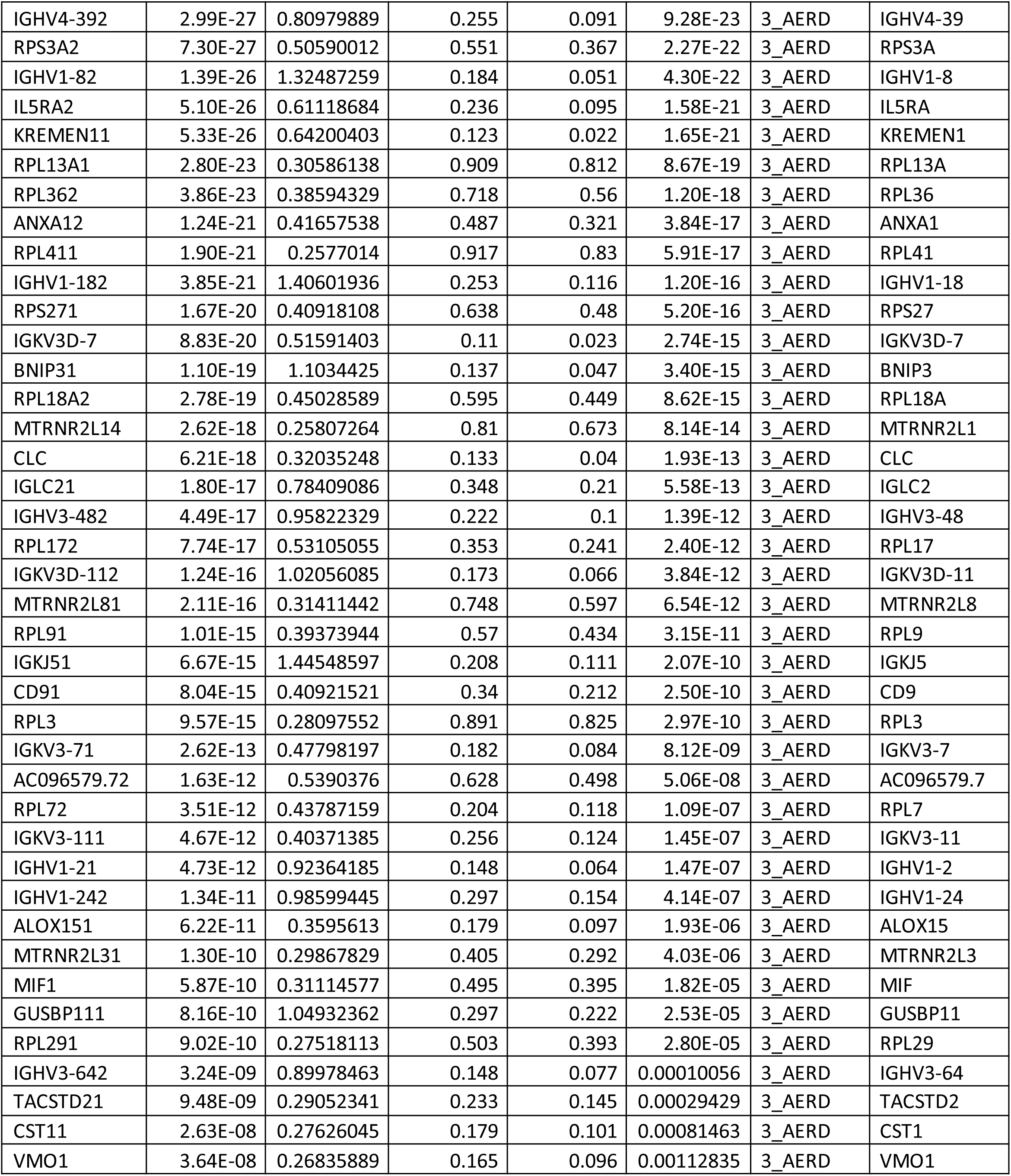
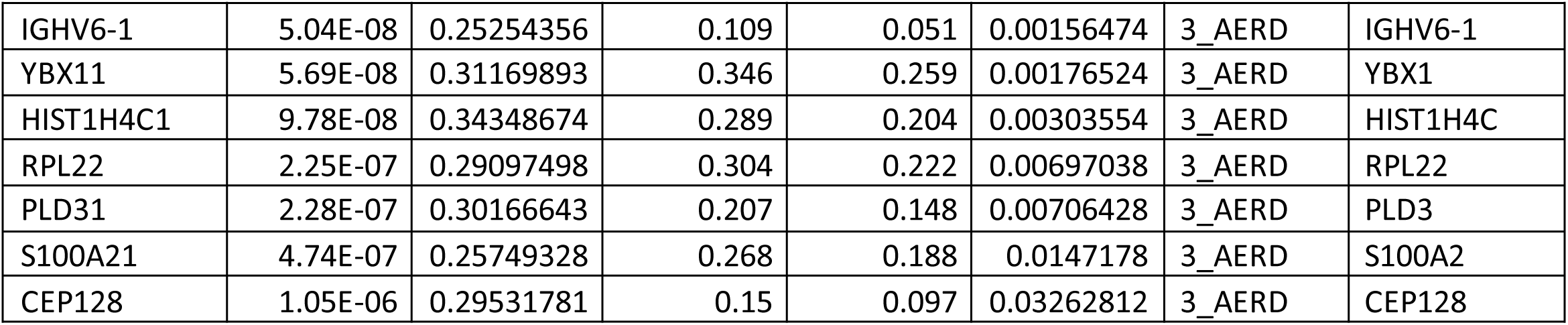
Patient-specific genes.

## Abbreviations

AERD: Aspirin-exacerbated respiratory disease
CRSwNP: Chronic rhinosinusitis with nasal polyps
CRSsNP: Chronic rhinosinusitis without nasal polyps
COX: Cyclooxygenase
Ig: Immunoglobulin
IL: Interleukin
LT: Leukotriene
PCA: Principal component analysis
PG: Prostaglandin
scRNA-seq: Single-cell RNA-sequencing
*UMAP*: Uniform Manifold Approximation and Projection

